# Sex, not estrous cycle stage, drives differences in the microglial transcriptome in the 5xFAD mouse model of amyloidosis

**DOI:** 10.1101/2025.03.28.646050

**Authors:** L. Calcines Rodriguez, N. Noyes-Martel, J.L Becker, A.K. Majewska, M.K. O’Banion

**Affiliations:** Department of Neuroscience, Del Monte Institute for Neuroscience, University of Rochester, Rochester, NY; Rochester Genomics Center, University of Rochester, Rochester NY; Center for Visual Science, University of Rochester, Rochester NY; Department of Neurology, University of Rochester, Rochester NY

**Keywords:** Microglia, Sex, Estrous, 5xFAD, Amyloid, Alzheimer’s, Transcriptome, Plaques

## Abstract

Alzheimer’s disease (AD) presents with a sex bias where women are at higher risk and exhibit worse cognitive decline and brain atrophy compared to men. Microglia play a significant role in the pathogenesis and progression of AD and have been shown to be sexually differentiated in health and disease. Whether microglia contribute to the sex differences in AD remains to be elucidated. Herein, we characterize the sex differences in amyloid-beta (Aβ) plaque pathology and microglia-plaque interaction using the 5xFAD mouse model of amyloidosis and further elucidate the microglial transcriptomic changes that occur in males and females. In females we concentrate on two hormonally distinct stages of the rodent estrous cycle: proestrus and diestrus. Our results indicate that Aβ plaque morphology is sexually distinct with females having greater plaque volume and lower plaque sphericity compared to males. Microglia also interact with plaques in a sexually distinct manner with females phagocytosing Aβ to a greater extent compared to males. Furthermore, we found that female microglia are not overtly different at the proestrus or diestrus stages. However, we found stark sex differences between female and male microglia transcriptomes in the 5xFAD brains, where female 5xFAD microglia were enriched in genes involved in glycolytic metabolism, antigen presentation, disease-associated microglia and microglia neurodegenerative phenotype (DAM/MGnD), and interferon signaling compared to male 5xFAD microglia.

## Introduction

Historically, researchers have neglected sex as a biological variable in study design, resulting in an incomplete understanding of disease pathogenesis and ultimately leading to misinformed clinical trials^1,2^. Alzheimer’s disease (AD) is a neurodegenerative disease that presents with a sex bias and is characterized by aberrant aggregation of the amyloid-β (Aβ) protein into senile plaques and the tau protein into neurofibrillary tangles^3–5^. This proteinopathy leads to synaptic and neuronal loss, microgliosis, astrogliosis, neuroinflammation, oxidative, and metabolic stress, ultimately resulting in cognitive decline^6^. Women with AD exhibit greater rates of cognitive decline, brain atrophy, and increased global pathology compared to age-matched men^3^, however, the intricate mechanisms behind these differences remain elusive, presenting challenges to the development of sex-specific therapeutic interventions.

Microglia, the resident immune cells of the central nervous system (CNS), play a variety of functions, including maintaining homeostasis through dynamic surveillance and phagocytosis of debris, with specific roles in neurogenesis, synaptic pruning, and plasticity^7–11^. In the context of AD, microglia are known to congregate around Aβ plaques^12,13^ and adopt unique transcriptional signatures including disease-associated microglia (DAM)^14^, activated response microglia (ARM)^15^, and microglial neurodegenerative phenotype (MGnD)^16^. Moreover, they are heavily involved in the phagocytosis, seeding, spreading, growth, and structural remodeling of Aβ plaques^17–23^. Their critical roles in AD pathogenesis were confirmed when genome-wide association studies revealed that many genetic risk factors for late-onset AD are almost exclusively or highly expressed in microglia in the CNS^24^. Microglia can also modulate the course of disease pathology by releasing anti- and pro-inflammatory cytokines and other secreted molecules that may ameliorate or aggravate pathological conditions^25–27^. Interestingly, these highly plastic cells are not only transcriptionally diverse and regionally heterogenous but also sexually differentiated in health and disease^28–36^.

Sex influences numerous biological processes and leads to significant divergence as a function of age, particularly in the brain, where cells are sexually differentiated from developmental stages to adulthood and beyond^36–38^. Sex differences can arise from the sex chromosomes (XX in females and XY in males) and gonadal hormones, both of which establish sex-specific features early in development through epigenetic modifications and activate sex-specific phenotypes later in life^36,37,39^. Apart from a developmental role, changes in gonadal hormones during the menstrual cycle in adult women have been linked to changes in cortical thickness in imaging studies^40,41^. In female rodents, fluctuations in spine density across the rodent estrous cycle are well documented^42^, and emerging evidence reveals that ventral hippocampal neurons shift their epigenetic landscape and transcriptomic profile at two hormonally distinct stages of the estrous cycle^43^.

It is postulated that microglia exhibit sex differences in their developmental trajectory and in adulthood, resulting in sex-specific variations in morphology, cell density, proliferation, migration, transcriptome, proteome, and overall function^31,34–37^. In the context of injury and disease, microglia can also respond in a sexually distinct manner^31,44–49^. For instance, in response to ischemic injury, female microglia minimize the damage more efficiently than males, and this ability seems to be sex-specific and cell intrinsic^44^. Currently, we do not fully appreciate the relative contributions of sex chromosomes and gonadal hormones to differences in microglia gene expression or function in health or disease. Understanding how biological sex influences microglia function may provide insights into the sex-specific prevalence of neurological and psychiatric diseases and allow for more personalized therapeutic interventions.

The present study seeks to investigate the sex differences in microglial responses to Aβ pathology and examine whether microglia gene expression varies at different stages of the estrous cycle. To this end, we characterized the differences in Aβ plaque morphology and microglia-plaque interaction in 5.5 moth old 5xFAD mice. We then isolated cortical microglia from 5.5-month-old wild type (WT) and 5xFAD mice of both sexes that develop extensive Aβ pathology and performed bulk mRNA sequencing (RNAseq). To further interrogate the effect of endogenous hormonal fluctuations on the microglial transcriptome, we selected females at proestrus and diestrus estrous stages, characterized by high and low levels of estrogen, respectively^50^. Our results indicate that plaque morphology is structurally distinct between female and male 5xFAD mice, and despite no difference in microglia-plaque coverage, female microglia seem to engulf more Aβ than male microglia. Our RNAseq data revealed sex-specific transcriptomic differences between female and male 5xFAD microglia, particularly in pathways related to glycolytic metabolism, type-I interferon signaling, and antigen presentation, as well as disease-associated phenotypes, with no significant differences noted in females at different stages of the estrous cycle.

## Methods

### Experimental animals

All animal procedures were reviewed and approved by the University Committee on Animal Resources of the University of Rochester Medical Center and performed according to the Institutional Animal Care and Use Committee and guidelines from the National Institute of Health (NIH). Animals were maintained on a 12:12h light:dark cycle with access to food and water ad libitum. Male and female 5xFAD mice (B6.Cg-Tg(APPSwFlLon,PSEN1*M146L*L286V)6799Vas/Mmjax, MMRRC stock 34840) were obtained from MMRC and maintained at the University of Rochester vivarium. These 5xFAD mice harbor Swedish, Florida and London mutations in the amyloid precursor protein (APP) and the M146L and L286V mutation in presenilin 1 (PSEN1). They begin accumulating plaques starting at 2 months of age and have a rapid and progressive increase in amyloid plaque load. Non-transgenic wild-type (WT) female and male littermates were used as controls where applicable. Mice were sacrificed at 5.5-6 months of age as 5xFAD mice exhibit extensive amyloid plaque pathology at these timepoints. Two weeks prior to sacrifice, vaginal smears were assessed daily (10.00 - 11.00 a.m.) in female mice. This equates to approximately 3 estrous cycles as females cycle over a period of 4-5 days through proestrus, estrus, metestrus, and diestrus stages^50^. At the time of brain collection, a final vaginal cytology assessment segregated the females into proestrus or diestrus. These stages were chosen as they exhibit high and low estrogen levels and mimic the late follicular and luteal phase of the menstrual cycle in humans, respectively^50^. Dissection of the uterus and subsequent weighing was used to confirm correct staging, as proestrus females tend to have bigger/heavier uterus compared to diestrus females. Only females who’s cycle staging and uterus weight were congruent were used for further analyses, including the RNA sequencing experiment.

### Vaginal cytology and uterus dissection

Estrous cycle staging was assessed based on vaginal smear and cytological examination (S.Fig.1). Two-weeks prior to sacrifice, a pipette tip filled with 40 µL of PBS was used to lavage the area around the vaginal orifice, avoiding any penetration. Samples were smeared onto a glass slide and dried. Slides were submerged in methanol for 30 s for fixing followed by a 1 min wash in eosin and then 30 s in hematoxylin. Slides were then washed with water and dried before examination under the microscope. Staging was based on the preponderance of nucleated vs. cornified squamous epithelial cells and leukocyte presence (S.Fig.1b). To meet the criteria for proestrus and diestrus, smears needed to show approximately 90% small nucleated epithelial cells or approximately 90% leukocytes, respectively. At the time of sacrifice, the uterus was dissected to exclude the ovaries and vaginal canal and subsequently weighed. Based on previous literature, uterine weight is greater in proestrus compared to diestrus stages^51^, as was replicated in our own data (S.Fig.1c, d).

**Figure 1.**
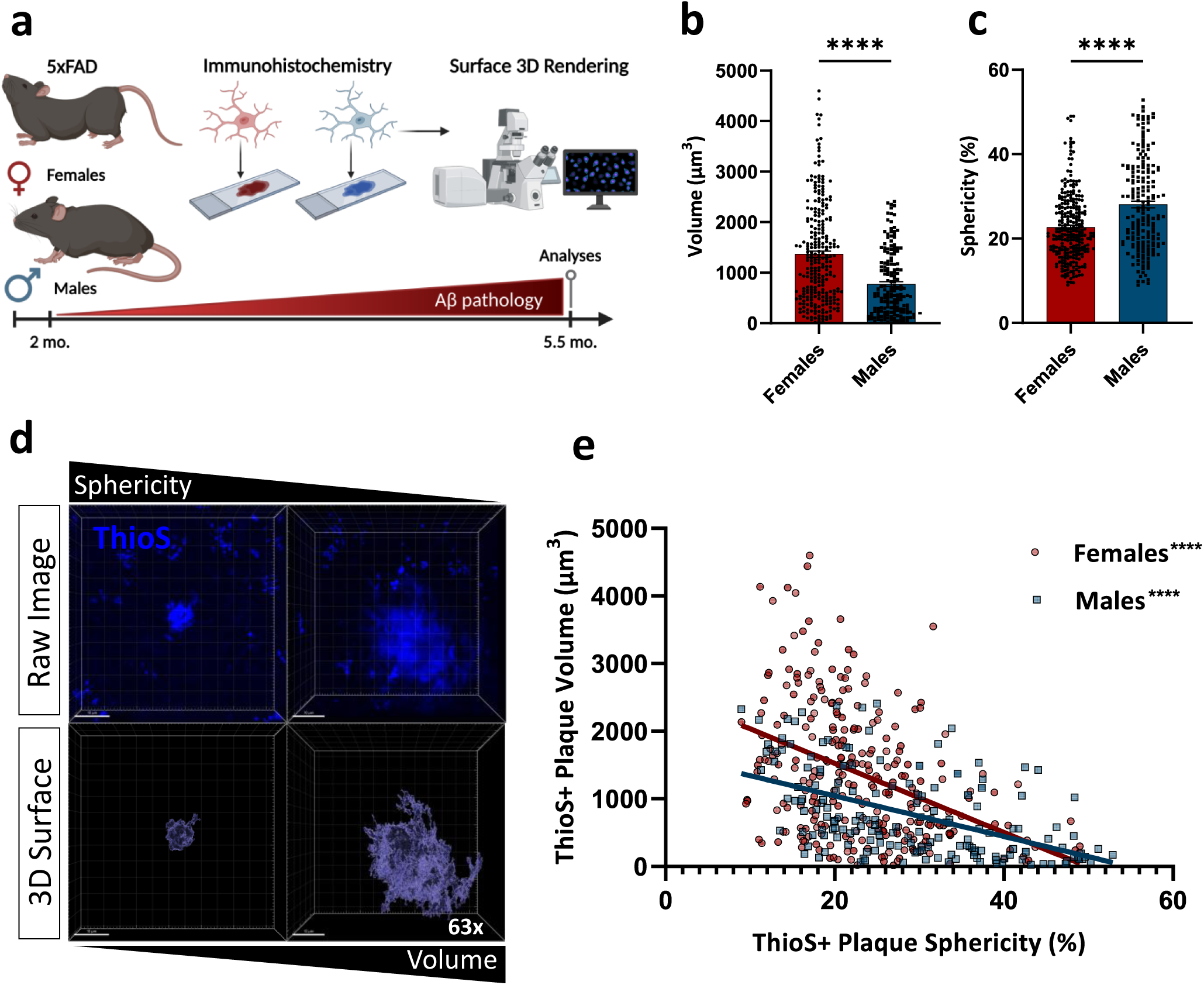
Amyloid-β plaque morphology is sexually differentiated in 5xFAD mice. a) Schematic of experimental design: Female and male 5xFAD mice were aged to 5.5 months and their brains used for immunohistochemistry followed by high resolution confocal imaging and 3D surface rendering. b) Quantification of cortical ThioS+ plaques volume (µm^3^) in female and male 5xFAD mice. Each data point represents an individual plaque analyzed at high magnification (63x magnification; n = 173-265 plaques/group) derived from three 5xFAD mice/group. Two-tailed Mann Whitney test, *\*\*\*\*p<0.0001*. Data is presented as mean ± SEM. c) Quantification of cortical ThioS+ fibrillar plaques sphericity (arbitrary units; AU) in female and male 5xFAD mice. Each data point represents an individual plaque analyzed at high magnification (63x magnification; n = 188-277 plaques/group) derived from three 5xFAD mice/group. Two-tailed Mann Whitney test, *\*\*\*\*p<0.0001*. Data is presented as mean ± SEM. d) Representative 63x confocal raw image (top) and 3D surface rendition (bottom) of ThioS+ fibrillar plaque with varying degrees of sphericity and volume (Scale bar = 10 µm). e) Correlation between ThioS+ plaque volume (µm^3^) and sphericity (%) in female and male 5xFAD mice. Two-tailed Spearman correlation; Female r = −0.4113, Male r = 0.5394; *\*\*\*\*p<0.0001*, Each data point represents an individual plaque analyzed at high magnification (63x magnification; n **=** 173-265 plaques/group) derived from three 5xFAD mice/group.

### Brain collection

Mice were injected with a mixture of xylazine (10 mg/kg, i.p.) and ketamine (100 mg/kg, i.p.) and perfused intracardially with 0.15 M phosphate buffered saline (PBS) containing 0.5% sodium nitrite and 2 IU heparin/mL. Following perfusion, the brain hemispheres were separated: one hemisphere was fixed in 4% paraformaldehyde (PFA, pH 7.2) for immunohistochemistry (described below) and the other hemisphere was processed for flow cytometry (described below).

### Immunofluorescence

Following perfusion, half-brains were fixed in 4% PFA overnight then transferred to 30% sucrose for 24 h. Brains were then flash frozen in dry ice and cryosectioned into 30 µm sections on a freezing-stage sliding microtome. Free-floating sections were stored in a cryoprotectant at - 20°C until assayed. Immunohistochemical assessment of proteins of interest involved the following steps: multiple washes with PBS to remove cyroprotectant, incubation in 4% normal goat serum (NGS) in 0.4% PBST (Triton X-100 (Sigma-Aldrich, #X100) diluted in PBS) for 1.5 h at room temperature (RT) to prevent non-specific binding and allow cell membrane permeabilization, respectively, followed by staining with primary antibodies with 4% NGS in 0.4% PBST for 72 h at 4°C. Next, the tissue was washed multiple times with PBS and incubated in secondary antibody in 4% NGS in 0.4% PBST for 1.5 h at 4°C. For Aβ staining, the tissue was then washed with PBS and incubated in 0.025% Thioflavin S (ThioS; Sigma-Aldrich, #T-1892) in water for 5 min at RT followed by multiple washes in 50% ethanol at RT. Slides were mounted on 2x subbed slides with Prolong Gold anti-fade mounting media (Invitrogen, #P36930) and cover slipped. Primary and secondary antibody dilutions included rabbit anti-IBA1, a pan-microglial marker, (Wako, #019-19741, 1:1000), biotinylated 6E10 (BioLengend, #803008, 1:3000), goat anti-rabbit Alexa 488 (Invitrogen, #A21245, 1:1500), and Streptavidin Alexa 594 (Invitrogen, #S32356, 1:1500).

### Image Acquisition and Analysis

For high resolution imaging, individual cortical plaques from 3 coronal brain sections (approximately 60-90 plaques/section) were imaged (n = 3 5xFAD brains per group) with a Leica Stellaris 5 Laser Scanning Confocal Microscope using a 63x oil objective (N.A. 1.4, HC PL APO 63x/1.4 OIL CS2, Leica Microsystems) and a zoom factor of 3.1. Images of the entire plaque were taken with a Z-step size of 0.3 µm, a pixel resolution of 1024 x 1024, and an 8-bit color depth. All parameters (gain and intensity) stayed consistent across samples. Bitplane IMARIS v10.2.0 was used for image analysis. IMARIS “surface” feature was used to detect IBA1+ cells as well as ThioS+ and 6E10+ plaques with a surface grain size of 0.58 µm. Image pre-processing for ThioS included a background subtraction based on the relative intensities of the measured background (*i.e.,* an area of the slide with no tissue) due to observable differences in intensity of the background, followed by a gaussian filter. Surface filters of >100µm^2^ and >130µm^2^ area were applied to avoid rendering small plaque seeds within the field of view for ThioS and 6E10, respectively. Pre-processing for IBA1 included a background subtraction and a surface filter of >800 voxels to avoid inclusion of small background speckles. The IMARIS batch processing feature was used across all samples and images were visually inspected to confirm 3D rendition closely resembled the raw image. For Fig.1 c, e and S.Fig.2d, sphericity was calculated by IMARIS using the following formula: ψ = (π^1/3^(6V_p_)^2/3^)/A_p_ where p is the particle (*i.e.,* the plaque). For Fig.2b, microglia plaque coverage was quantified by dilating the surface of ThioS+ plaques by 2 µm and measuring the overlapping voxels between the dilated ThioS surface and the IBA1 surface. The resulting value was then normalized to the surface area of the ThioS+ plaque. For Fig.2c-e, IBA1 and ThioS overlap was obtained by quantifying the overlapping voxels between the IBA1 and ThioS surface and normalizing it to ThioS total volume. For Fig.2c, plaques were categorized into low, medium, and high sphericity using the equal width binning method. For global plaque density analysis, 3 coronal brains sections were imaged (n = 4-6 5xFAD brains per group) using a 10x objective (N.A. 0.4, HC PL APO 10x/0.40 CS2, Leica Microsystems) and a digital zoom factor of 2.25. Images of the entire section were taken with a Z-step size of 2.41 µm, a pixel resolution of 512 x 512, and an 8-bit color depth. All parameters (gain and intensity) stayed consistent across samples. IMARIS “Spots” feature was used to quantify the number of plaques within a pre-specified ROI (cortex). The number of spots (*i.e.,* number of plaques) was normalized to the area of the ROI yielding global plaque density (plaque #/µm^2^). Experimenter was blinded to the sex of the 5xFAD mice until after the end of data collection/IMARIS analysis.

**Figure 2.**
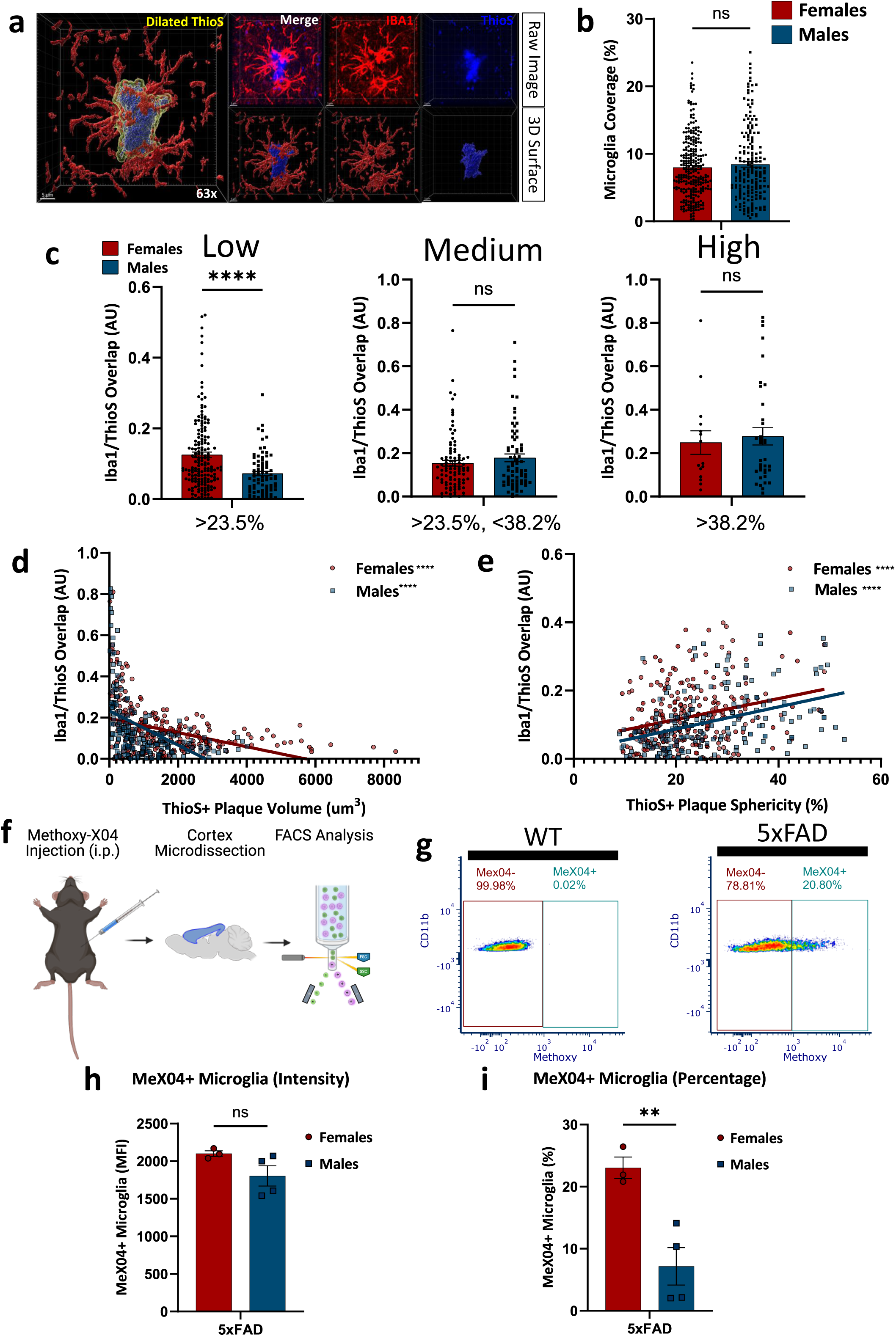
Microglia-plaque interaction is sexually differentiated in the 5xFAD mice. a) Representative 63x confocal image and 3D surface rendition of Iba1 (microglia; red), ThioS (Aβ plaque; blue) and the 2 µm-dilated surface of ThioS (yellow) to assess microglia plaque coverage of the plaque (Scale bar = 5 µm). b) Quantification of the percent of microglia plaque coverage around ThioS+ plaques in female and male 5xFAD mice. n = 185-270 plaques/group derived from three 5xFAD mice/group. Two-tailed Mann Whitney test. Data is presented as mean ± SEM. c) Quantification of Iba1/ThioS+ overlap (normalized to ThioS volume; AU) in low (n = 78-166 plaques/group; includes plaques with <23.5% sphericity), medium (n = 74-96 plaques/group; includes plaques with >23.5 and <38.2% sphericity), and high (n = 15-36 plaques/group; includes plaques with <38.2% sphericity) sphericity plaques, as determined by equal-width binning method. Each data point represents an individual plaque analyzed at high magnification and derived from three 5xFAD mice/group. Two-tailed Mann Whitney test, *\*\*\*\*p<0.0001*. Data is presented as mean ± SEM. d) Correlation between IBA1/ThioS+ overlap (normalized to ThioS volume; AU) and ThioS plaque volume (µm^3^) in female and male 5xFAD mice. Two-tailed Spearman correlation; Female r = −0.3995, Male r = −0.5933; *\*\*\*\*p<0.0001*, Each data point represents an individual plaque analyzed at high magnification (n **=** 188-277 plaques/group) derived from three 5xFAD mice/group. e) Correlation between Iba1/ThioS+ overlap (normalized to ThioS volume; AU) and ThioS plaque sphericity (%) in female and male 5xFAD mice. Two-tailed Spearman correlation; Female r = 0.2536, Male r = 0.4378*; ****p<0.0001*, Each data point represents an individual plaque analyzed at high magnification (n **=** 171-264 plaques/group) derived from three 5xFAD mice/group. f) Schematic of experimental design: Female and male 5xFAD mice of 6 months of age were intraperitoneally (i.p.) injected with Methoxy-X04 dye (MeX04; 4 mg/kg) 24 h prior to cortex microdissection for subsequent flow cytometry analysis. g) Representative flow cytometry dot plots showing cells that were Cd11b^+^MeX04^-^ and Cd11b^+^MeX04^+^ after identification of live microglia (7AAD^-^Cd11b^+^CD45^Low-Intermediate^; not shown) in MeX04-injected WT and 5xFAD mice. h) Quantification of the mean fluorescence intensity (MFI) of the MeX04 signal within MeX04^+^ microglia in female and male 5xFAD mice (n = 3-4 mice/group). Two-tailed Unpaired t test. Data is presented as mean ± SEM. i) Quantification of the percentage of MeX04^+^ microglia in female and male 5xFAD mice (n = 3-4 mice/group). Two-tailed Unpaired t-test. Data is presented as mean ± SEM.

### Methoxy-04 injection and flow cytometry/Fluorescent Activated Cell Sorting (FACS)

Mice were intraperitoneally (i.p.) injected with 4 mg/kg of Methoxy-04 (Tocris Bioscience, #4920) 24 h prior to sacrifice and brain collection as previously described. The cortex was microdissected from the half brain and tissue was Dounce homogenized in 3 mL of FACS buffer (0.5% BSA in 1X PBS). Brain homogenates were then filtered through a 70 µm filter (MACS, #130-100-916) into a 15 mL canonical tube containing 3 mL of FACS buffer, rinsed with 3 mL additional FACS buffer, and centrifuged at 600 g for 6 min at 4°C. Cell pellet was then resuspended in 9 mL of 40% Percoll (Cytiva, #17089101) diluted in 1X PBS and centrifuged at 2000 g for 20 min at 4°C with the break off to avoid disrupting the gradient. Myelin and supernatant were removed and cell pellet was resuspended and transferred to a new tube containing 8 mL of FACS buffer followed by centrifugation at 600 g for 6 min at 4°C. Supernatant was discarded and cell pellet was transferred into a new 1.5 mL Eppendorf tube, resuspended in 90 uL of FACS buffer containing Fc block (BD Pharmingen, #553141, 1:90), and incubated for 15 min at 4°C. Next, 10 uL of antibody master mix containing Cd11b-PerCP/Cy5.5 (BD Biosciences, Clone M1/70, 1:400) and CD45-APC/Cy7 (BioLegend, Clone 30-F11, 1:400) was added to the cells and incubated for 30 min in the dark at 4°C. Following incubation, cells were washed with FACS buffer and centrifuged at 400 g for 5 min at 4°C. Cells were then transferred into 5 mL Falcon tubes containing 7AAD Viability Staining Solution (eBioscience, #00-699-50, 1:80). Compensation (*i.e.,* single-stain) and fluorescence-minus-one (FMO) controls were run in-tandem with samples using compensation beads (Invitrogen, #01-2222-41) or cells which informed the gating criteria. Samples were run on a FACSAria II (BD) cytometer. Following debris, doublet, and dead cell exclusion, microglia were identified as Cd11b+/CD45^low-intermediate^ and were further divided into a Methoxy-04 positive or negative population. Data was analyzed using FCS express 7 (DeNovo Software, v7.16.0035) and geometric mean fluorescence intensity (MFI) along with the percentage of cells within each gate was calculated for each population after compensation.

### Microglia isolation and bulk mRNA sequencing

Microglia were isolated from the microdissected cortex and processed for FACS as described above. This time, samples were instead incubated with 10 uL of antibody master mix containing Cd11b-FITC (BioLegend, Clone M1/70, 1:400) and CD45-APC/Cy7 (BioLegend, Clone 30-F11, 1:400), TMEM119-PE (Abcam, Clone 106-6, 1:500) and P2RY12-APC (BioLegend, Clone S16007D, 1:50) and subsequently stained with 7AAD Viability Staining Solution (eBioscience, #00-699-50, 1:80). Compensation and FMO controls were included, and samples were run on an Aurora CS (Cytek) spectral cytometer. Following debris, doublet, and dead cell exclusion, microglia were identified as Cd11b+/CD45^low-intermediate^ and TMEM119 and P2RY12 were used to further confirm microglial identity. Cells were sorted into RLT Plus (Qiagen) + 10 uL/mL β-mercaptoethanol (BME) and stored at −80°C until processing for bulk RNAseq.

Total RNA was isolated using the RNeasy Plus Micro Kit (Qiagen, Valencia, CA). The concentration of RNA was determined using the NanoDrop One spectrophotometer (NanoDrop, Wilmington, DE) and RNA quality was assessed with the Agilent Bioanalyzer 2100 (Agilent, Santa Clara, CA). Pre-amplification of 1 ng of total RNA was performed with the SMART-seq v4 Ultra-Low RNA Input kit (Clontech, Mountain View, CA) per manufacturer’s instructions. Quantity and quality of the cDNA was determined using Qubit Fluorometer (Life Technologies, Carlsbad, CA) and the Agilent Bioanalyzer 2100 (Agilent, Santa Clara, CA). A total of 150 pg of cDNA was used to generate Illumina compatible sequencing libraries with the NexteraXT library preparation kit (Illumina, San Diego, CA) per manufacturer’s instructions. The libraries were then hybridized to the Illumina flow cell and sequenced using the 10B-300 NovaSeq X sequencer (Illumina, San Diego, CA) to obtain approximately 30 million paired-end reads of 150 nucleotides for each sample. Four biological replicates were sequenced for each group.

### Bioinformatic analyses

Following sequencing, the resulting RNAseq raw reads were demultiplexed using bcl2fastq v2.20.0 followed by quality filtering and adapter removal with FastP (v0.23.1). Cleaned reads (FASTQ files) were then mapped onto the GRCm39/gencode M31 reference using STAR (v2.7.9a), generating BAM output files. Gene-level read quantification was derived with subread (v2.0.1) package (featureCounts) with a GTF annotation file (GRCm39/gencode M31). Differential expression analysis was performed using DESeq2 (v1.44.0) with an adjusted p-value (FDR) threshold of 0.05 and LFC cutoff of ±0.25 within R (v4.4.1, https://www.R-project.org/). A PCAplot was created within R using the ‘pcaExplorer’ (v2.30.0) to measure the variance of gene expression (top 500 genes) in each sample. Heatmaps were generated using the ‘pheatmap’ package (v1.0.12) given the Z-normalized rLog transformed expression values. Gene ontology analyses were performed with the EnrichR package (v3.2). Volcano plots and dot plots were created using ggplot2 (v3.5.1). Venn diagrams and resulting list of overlapped or unique genes were derived from FunRich (v3.1.3) and plotted using BioRender. The following list of input genes were used in Venn diagrams: canonical glycolysis (Alliance Genome Consortium orthologs based on Hallmark Glycolysis, v2024.1.Mm), XY genes (GenBank, Mus musculus strain C57BL/6J chromosome X and Y, GRCm39), antigen presentation (Antigen Processing and Presentation, GO:0019882, v2024.1.Mm), DAM genes^14^, MGnD genes^16^.

### RT-qPCR

Flash frozen cortices from 5.5-month-old 5xFAD mice were homogenized using a tissue homogenizer (Omni International) in RLT Plus Buffer with 10 uL/mL of BME. RNA was extracted using the RNeasy Plus Mini kit (Qiagen) as per manufacturer’s instructions. cDNA was acquired from 1 ug of RNA using TranscriptMe RNA kit (Blirt, Qiagen, RT31-020). Concentration of RNA and cDNA was measured using NanoDrop 2000 (ThermoFisher Scientific). Real-time quantitative polymerase chain reaction (RT-qPCR) was run using a 10 uL reaction in a 96-well plate on QuantStudio Real Time PCR System (ThermoFisher Scientific). Two separate reactions were assembled using 100 ng of cDNA, TaqMan Fast Advanced Master Mix (Applied Biosystems, 2X) and the following TaqMan Gene Expression Assays: human APP (Hs00169098_m1, FAM-MGB, 20X), human PSEN1 (Hs00997789_m1, FAM-MGB, 20X), and murine GAPDH (Mm99999915_g1, VIC-MGB, 20X) as a reference control. A total of 6 biological replicates (cortices from individual 5xFAD mice) were used with 4 technical replicates per sample. Fold changes were quantified using the 2^-ΔΔCt^ method and experimenter remained blind to the sex of the samples up to the moment of data analysis.

### Statistical analyses

All statistical analyses were performed in Prism (GraphPad Software, v10.4.1, San Diego, CA), except for the RNAseq analyses which were performed in R as described under “Bioinformatic analyses”. Depending on the distribution of the data, as tested using the Shapiro-Wilk test, parametric and non-parametric tests were performed including unpaired student’s t test, Mann Whitney test, two-way ANOVA, Spearman correlation, and simple linear regression. Outliers were identified and removed using the Robust Regression and Outlier Removal (ROUT) and Grubb’s method, removing multiple or one outlier, respectively. The following number of outliers were removed using the ROUT method (Q = 1%): 27 (Fig.1b), 27 pairs (Fig.1e), 40 (S.Fig.2c), 6 (Fig.2b), 30 pairs (Fig.2d). The following outliers were removed using the Grubbs method: 1 (S.Fig.2b), 1 (Fig.2h), 1 (Fig.2i).

## Results

### Plaque morphology is sexually differentiated in female and male 5xFAD mice

Previous studies have reported sex differences in global Aβ plaque burden in the 5xFAD mouse model^52^. Here, we sought to confirm whether cortical Aβ plaque morphology differed between the sexes, given that Aβ morphology, including Aβ plaque volume and compaction, have been shown to be modulated by microglia and can impact neuronal health (*e.g*., neuritic dystrophy)^12,13,18,19^. To this end, we examined Aβ plaque morphology in 5xFAD female and male mice at 5.5 months of age, a time when the pathology is well manifested in this murine mouse model of amyloidosis (Fig.1). We utilized immunohistochemistry combined with high-resolution imaging to generate 3D surface renderings of individual cortical plaques in each sex (Fig.1a). To examine Aβ plaques, we used ThioS which binds to fibrillar β-sheet conformations of Aβ and 6E10 which recognizes all conformations. When assayed at high magnification (63x), we found that both ThioS+ (Fig.1b) and 6E10+ (S.Fig.2c) individual plaque volume was significantly greater in the cortex of female 5xFAD mice compared to males. When assessing global 6E10+ plaque density in the entire cortical region, we noted that females had greater plaque density compared to males (S.Fig.2b). Importantly, when measuring the mRNA levels of the human *APP* and *PSEN1* transgenes in the 5xFAD brain, we did not find any statistically significant sex differences (S.Fig.2e, f), suggesting that the sex differences in Aβ load are not due to differences in transgene expression. Next, we sought to investigate plaque sphericity which is often used as a proxy for plaque compaction^53,54^. We observed that cortical ThioS+ plaque sphericity was significantly lower in female 5xFAD mice compared to males (Fig.1c), suggesting that female cortical fibrillar plaques are, not only larger, but less compact (Fig.1b-d). Indeed, there was a significant negative correlation between plaque volume and sphericity in both sexes (Fig.1e) reinforcing the notion that larger plaques are less compact. Interestingly, this sex difference in plaque sphericity was not noted in 6E10+ plaques (S.Fig.2d), suggesting that this morphological feature is specific to fibrillar forms of Aβ (*i.e.,* ThioS+ plaques). Taken together, these results reveal sex-specific differences in plaque morphology in 5xFAD mice, with females exhibiting a more pronounced Aβ burden.

### Microglia exhibit sexually differentiated plaque interaction in female and male 5xFAD mice

Given the observed sex differences in plaque morphology (Fig.1) and prior work implicating microglia in plaque compaction^12,13^, we sought to understand whether microglia remodel plaque morphology in a sex-specific manner in the 5xFAD mouse model. Owing to the close proximity of microglia to plaques, we assessed the extent of plaque coverage by IBA1, a pan-microglial marker in 5.5-month-old female and male 5xFAD brains (Fig.2a). To do this, we computed the overlap between a dilated ThioS surface and the IBA1 surface (Fig.2a), normalized to the surface area of the ThioS+ plaque to obtain a percentage of microglia-plaque coverage (Fig.2a, b). Surprisingly, we did not find a significant sex difference in microglia coverage of ThioS+ plaques between female and male 5xFAD mice (Fig.2b), contrasting a previous report that used a 5xFAD-ApoE3 mouse model^54^.

As brain-resident immune cells, microglia not only closely interact with plaques but also phagocytose Aβ species^55^, a process that has been shown to be involved in degradation as well as deposition and growth of Aβ plaques^19,20,56^. To assess whether there was a sex difference in microglial phagocytosis of Aβ, we examined the level of co-localized ThioS+ Aβ within IBA1^+^ cells. To this end, we computed the overlap between ThioS and IBA1 surfaces normalized to ThioS volume (Fig.2c-e). We found that females exhibited a significantly greater extent of ThioS+ plaque encapsulation by IBA1^+^ cells around plaques with low sphericity (Fig.2c). Correlation analyses revealed a significant positive relationship between co-localized IBA1 and ThioS vs. plaque sphericity in both sexes (Fig.2e). We also observed a significant negative correlation between co-localized IBA1 and ThioS vs. plaque volume in both sexes (Fig.2d). Once again, these correlations highlight the relationship between plaque volume and sphericity.

Next, to validate whether microglia phagocytose Aβ in a sexually-differentiated manner, 6-month-old female and male 5xFAD mice were injected 24 h prior to brain collection with Methoxy-X04 (MeX04), a blood-brain barrier penetrant dye that labels Aβ fibrils^57,58^ (Fig.2f). We subsequently microdissected the cortex and used flow cytometry to identify microglia (Cd11b^+^CD45^low-intermediate^ cells) that were positive or negative for MeX04 (Fig.2g). When assessing the mean fluorescence intensity (MFI) of the MeX04 signal within each microglia, we did not find a statistically significant difference between microglia from the two sexes (Fig.2h). However, we did observe that female 5xFAD mice had greater numbers of MeX04+ microglia compared to males (Fig.2i), consistent with our histological data. Taken together, these results suggest that microglia interact with plaques in a sex-specific manner.

### Transcriptomic analysis reveals no overt differences in cortical microglia from 5xFAD females at different stages of the estrous cycle

Having found that plaque morphology is sexually distinct and that microglia physically interact with plaques in a sex-specific manner, we next wanted to assess the microglial transcriptomic profile in response to cortical Aβ plaques in female and male 5xFAD mice (Fig.3). Although our histological work only compared female and male 5xFAD mice, we elected to carry out our transcriptional analysis using females at different stages of the estrous cycle. To our knowledge, no other study has investigated whether microglia could present different phenotypes depending on estrous cycle stage. For this reason, we used vaginal cytology (S.Fig.1b) and uterine weight assessment to group the females into proestrus or diestrus stages (S.Fig.1c, d). Our selection is based on the well-established differences in the hormonal milieu at these stages, with the estrogen to progesterone ratio being high in proestrus and low in diestrus^50,59^. We found that both WT and 5xFAD females cycled regularly and exhibited similar changes in uterine weight, with proestrus females having approximately a 2-fold increase compared to diestrus females (S.Fig.1c, d), consistent with previous reports^51^.

**Figure 3.**
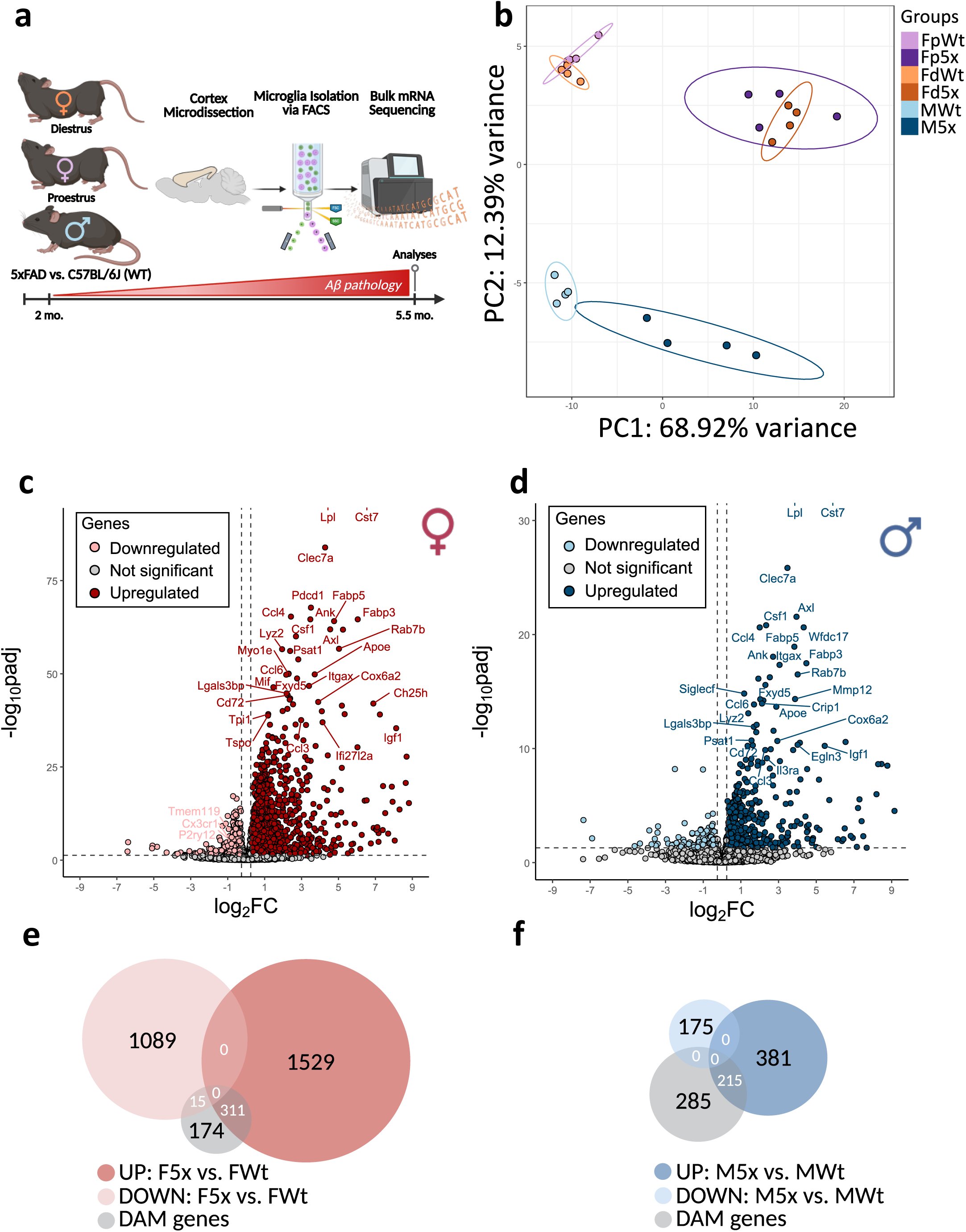
Microglia respond to amyloid-β in both sexes, with females exhibiting a more robust DAM phenotype than males. a) Schematic of experimental design: Proestrus female, diestrus female, and male WT and 5xFAD mice were aged to 5.5 months and their cortex was microdissected for microglia isolation via FACS and subsequent processing for bulk mRNA sequencing. b) Principal component analysis (PCA) of the top 500 variable genes in each sample. (Fp = proestrus female, Fd = diestrus female, M = male) showing the first (PC1) and second (PC2) principal components. c-d) Volcano plot showing the DEGs in the pairwise comparison between 5xFAD and WT female (c) and male (d) microglia. *FDR<0.05; |LFC|>0.25*. e-f) Venn diagram of the resulting DEGs from the volcano plots (in c and d) that overlap with previously reported DAM genes in female (e) and male (f) microglia. F5x = female 5xFAD microglia, FWt = female WT microglia; M5x = male 5xFAD microglia, MWt = male WT microglia; UP = upregulated genes, DOWN = downregulated genes.

Following estrous cycle staging, bulk mRNA sequencing (RNAseq) was performed on isolated cortical microglia from 5.5-month-old WT and 5xFAD male, proestrus, and diestrus females (Fig.3a). Principal component analysis (PCA) revealed a clear separation between groups with the variance explained primarily by genotype (5xFAD vs. WT; PC1: 68.92%) followed by sex (Female vs. Male; PC2: 12.39%) (Fig.3b). Both the PCA (Fig.3b) and the Euclidian sample-to-sample distance heatmap (S.Fig.3a) revealed that all groups clustered within their respective genotype and sex, confirming there was little within-group but large between-group variability. Notably, proestrus and diestrus females clustered together within their genotype in the PCA space (Fig.3b) and indiscriminately in the Euclidian heatmap within their genotype (S.Fig.3a). Both results are indicative that the transcriptomic profiles of proestrus and diestrus female microglia do not significantly differ in either genotype. Indeed, heatmaps of the differentially expressed genes (DEGs) in a pairwise comparison revealed that proestrus and diestrus female microglia exhibited modest changes in gene expression in both the WT (S.Fig.3b) and the 5xFAD brain (S.Fig.3c). In WT mice, we observed 7 upregulated and 7 downregulated genes in diestrus microglia compared to proestrus microglia (S.Fig.3b). In 5xFAD mice, there were 7 upregulated and 5 downregulated genes in diestrus microglia compared to proestrus microglia (S.Fig.3c). Interestingly, *N4bp3* expression was upregulated in the 5xFAD proestrus female microglia (S.Fig.3e). *N4bp3* is involved in antiviral responses through the RIG-I-Like Receptor and Mitochondrial Antiviral Signaling protein (MAVS), which induces type I interferon (IFN) signaling^60^. Furthermore, proestrus female microglia also showed increased *Nlrp3* in both WT and 5xFAD mice (S.Fig.3d, e). *Nlrp3* is involved in the formation of the inflammasome, a structure required for production of cytokines such as interleukin-1β (IL-1β) and IL-18^61^. This suggests that proestrus female microglia may be more prone to inflammatory signaling compared to their diestrus female counterparts. However, due to the few DEGs identified, we conclude that estrous cycle stage does not appear to overtly affect microglial gene expression in the healthy or Aβ-diseased brain. Intriguingly, when compared to male 5xFAD microglia, there are distinct gene sets and functional pathways that are more pronounced in either proestrus or in diestrus 5xFAD female microglia (S.Fig.5), although these genes are not statistically significant in the direct pairwise comparison between proestrus and diestrus female 5xFAD microglia. Taken together and given that proestrus and diestrus female microglia are not overtly different in either WT or 5xFAD mice, we decided to group proestrus and diestrus females into a single female group within their respective genotypes for the remaining analyses.

### Female microglia show a more robust response to Aβ pathology, while male microglia retain a more homeostatic phenotype

Microglia are known to adopt distinct transcriptomic signatures in response to Aβ, namely DAM and MGnD, that are characterized by the downregulation of homeostatic microglia markers and upregulation of genes involved in lipid metabolism and phagocytosis^14,16^. These signatures are especially prominent in plaque-associated microglia and phagocytic microglia observed in the context of neurodegenerative diseases, including AD^14,16,62^. To confirm that microglia were responding to Aβ in both sexes, we performed pairwise comparisons between 5xFAD and WT female (Fig.3c) and male (Fig.3d) microglia. As expected, volcano plots revealed that both female and male 5xFAD microglia responded to Aβ pathology and upregulated DAM genes such as *Cst7, Lpl, Trem2, Tyrobp, Axl, Clec7a, Apoe, and Cstd*, amongst others (Fig.3c, d; Table S1). However, female 5xFAD microglia seemed to have approximately 3 times more total DEGs (2,944 DEGs in females: 5xFAD vs. WT; Fig.3e) than male 5xFAD microglia (771 DEGs in males: 5xFAD vs. WT; Fig.3f) when compared to their WT counterparts. Using Venn diagrams to examine the extent of overlap between the DEGs and previously reported DAM genes^14^, we found that when compared to their WT counterparts, female 5xFAD microglia had 65.2% differentially expressed DAM genes (326 out of 500 DAM genes; Fig.3e), whereas males had 43% (215 out of 500 DAM genes; Fig.3f). More specifically, female 5xFAD microglia had 311 upregulated genes that were shared with the DAM phenotype, including *Cst7*, *Lpl*, *Clec7a*, *Itgax*, *Spp1*, *Apoe*, *Axl*, and *Tyrobp*, and 15 downregulated genes, including canonical homeostatic makers such as *P2ry12*, *Tmem119*, and *Cx3cr1* compared to female WT microglia (Fig.3e; Table S1). On the other hand, male 5xFAD microglia had 215 upregulated genes that shared the DAM phenotype, including those mentioned for females; however, male 5xFAD microglia did not downregulate any genes associated with the DAM phenotype compared to male WT microglia (Fig.3f; Table S1). This suggests that microglia from both sexes are responding to Aβ pathology; however, females exhibit a more robust response, particularly enriched in the DAM phenotype, whereas males retain a more homeostatic signature.

### Female and male 5xFAD microglia exhibit sexually differentiated gene expression profiles

Microglia are known to show sex-specific functional responses to insults and in disease contexts^34,35,38,44^. To investigate whether there is a sex difference in cortical microglia in the healthy brain and in response to Aβ, we performed pairwise comparisons between female and male WT (Fig.4a) and 5xFAD (Fig.4c) microglia. Surprisingly, we did not find extensive differences between female and male WT microglia, except for well-known sex chromosome-linked genes^63,64^ (X linked: *Xist*, *Tsix*, *Kdm6a*, *Eif2s3x*, *Kdm5c*; Y-linked: *Uty*, *Kdm5d*, *Eif2s3y*, *Ddx3y*, and *Gm4017*) and a few autosomal genes (Fig.4a, b). However, we did find numerous genes that were differentially regulated between female and male 5xFAD cortical microglia (Fig.4c). Sex chromosome-linked genes included all the aforementioned genes, as well as a few additional genes, including *Cybb* and *Pgk1* (Fig.4d), which are involved in oxidative metabolism and glycolysis, respectively. The remaining DEGs between female and male WT and 5xFAD microglia were autosomal genes. For instance, the pairwise comparison between female and male 5xFAD microglia revealed that 93.2% of DEGs were autosomal genes (409 autosomal genes out of 439 total DEGs; Fig.4d). In the pairwise comparison between female and male WT microglia only 69.7% DEGs were autosomal genes (23 autosomal genes out of 33 total DEGs; Fig.4b). This is indicative that a chronic stimulus such as Aβ pathology induces a context-specific sex divergence that is not observed at baseline (*i.e.,* in the non-diseased brain) and extends beyond sex chromosome-linked genes. Interestingly, a hierarchically-clustered heatmap of all the DEGs resulting from the female and male 5xFAD microglia comparison shows that female 5xFAD microglia are distinct from all other groups (Fig.4e). For instance, it shows that male 5xFAD microglia and male WT microglia cluster indiscriminately, whereas female 5xFAD microglia cluster separately from female WT microglia and all other groups (Fig.4e). This further supports the notion that male 5xFAD microglia retain a more homeostatic phenotype, as they cluster with WT male microglia, whereas female 5xFAD microglia respond in a discrete manner.

**Figure 4.**
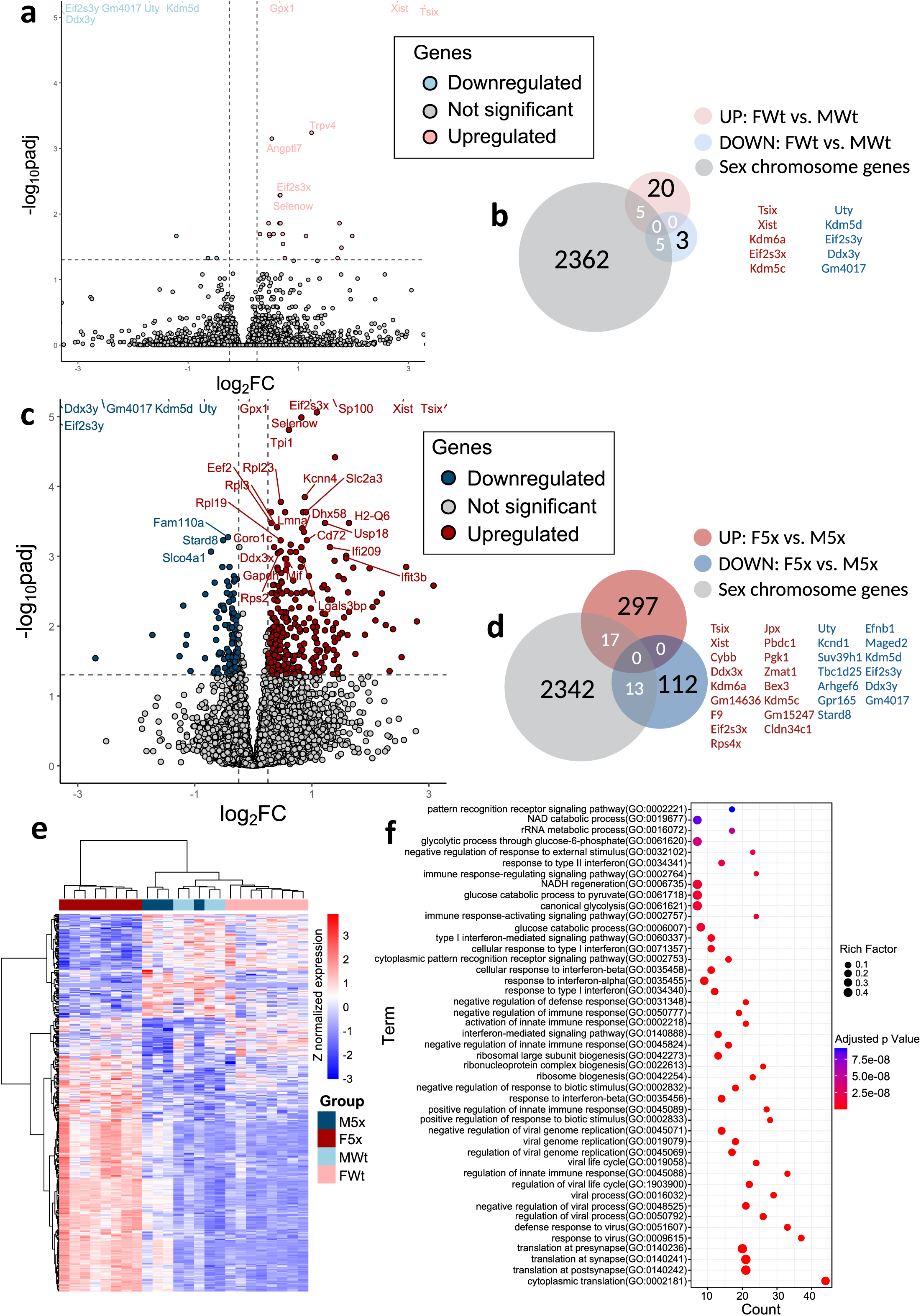
Microglia exhibit sex differences in autosomal gene and sex chromosome-linked gene expression in response to amyloid-β. a, c) Volcano plot showing DEGs in the pairwise comparison between female and male WT (a) and 5xFAD (c) microglia. *FDR<0.05; |LFC|>0.25*. b, d) Venn diagram of the resulting DEGs from the volcano plots (in a and c) that overlap with the 2372 known sex chromosome-linked genes (C57BL/6J X and Y chromosome, GRCm39) e) Heatmap with hierarchical clustering of all the DEGs (Z normalized) from the pairwise comparison between female and male 5xFAD microglia shown across all genotypes. M5x = male 5xFAD microglia, F5x = female 5xFAD microglia, MWt = male WT microglia, FWt = female WT microglia. f) Gene ontology analysis of the DEGs from the pairwise comparison between female and male 5xFAD microglia.

To query the functional roles of these DEGs, we performed Gene Ontology (GO) analysis with particular emphasis on biological process. The top upregulated pathways in female 5xFAD microglia compared to their male counterparts included pathways involved in glycolytic metabolism (*e.g.,* “glycolytic process through glucose-6-phosphate”, “canonical glycolysis”, “glucose catabolic process”, NADH regeneration”, NAD catabolic process”) (Fig.4f; Table S3). Glycolytic metabolism is enhanced in the context of AD, and microglia have been shown to undergo metabolic reprogramming in response to Aβ pathology^65,66^. This shift from oxidative phosphorylation to glycolytic metabolism for ATP production may enable microglia to carry out highly energy-demand processes^65^. Indeed, it has been reported that glycolytic microglia have enhanced phagocytosis, a process that requires rapid generation of ATP. Interestingly, the genes that were upregulated in female 5xFAD microglia compared to male counterparts included *Gapdh*, *Tpi*, *Pkm*, *Pgk1*, *Eno1*, *Aldoa*, and *Ldha*, all of which are glycolytic enzymes (S.Fig.4a, b). With the exception of *Mpi*, which was downregulated, other upregulated genes in female 5xFAD microglia that have been linked to canonical glycolysis included anabolic/catabolic enzymes such as *Got1*, *Pgam1*, as well as *Sdc3*, *Cdk1*, *Mif*, *Isg20*, and *Stmn1* (S. Fig.4a, b), the latter being involved in microtubule dynamics^67^.

Another set of pathways enriched in female compared to male 5xFAD microglia related to ribosomal biogenesis (“ribosomal large subunit biogenesis”, “ribonucleoprotein complex biogenesis”, “ribosome biogenesis”, “cytoplasmic translation”) amongst others (Fig.4f, Table S3). Surprisingly, we did not find canonical DAM genes (*Trem2*, *Ctsd*; S.Fig.4c) to be differentially expressed in female and male 5xFAD microglia. Instead, the majority of DAM genes found in this pairwise comparison were genes involved in ribosome biogenesis (S.Fig.4d, e). These included genes that encode for the ribosomal protein small subunit (*Rpsa*, *Rps4x*, *Rps8*, *Rps11*, *Rps18*, *Rps20*, *Rps27a*) and the ribosomal protein large subunit (*Rpl0*, *Rpl4*, *Rpl5*, *Rpl6*, *Rpl7*, *Rpl10a*, *Rpl11*, *Rpl12*, *Rpl14*, *Rpl18a*, *Rpl22*, *Rpl23*, *Rpl27a*) (S. Fig.5e; Table S2). Other genes within the DAM phenotype included glycolytic genes (*Pgk1*, *Ldha*, *Tpi1*, *Mif*, *Aldoa*, *Got1*, *Pgam1*) as well as antigen presentation genes (*H2-D1*, *H2-K1*, *Fgl2*) (S.Fig.4d, e; Table S2). *Apoe*, a gene that is a major genetic risk factor for AD and is shared amongst the DAM and MGnD phenotypes, was upregulated in female 5xFAD microglia compared to males (S.Fig.4d-g). Similarly, other MGnD genes, some of which overlap with DAM, were upregulated (*Cxcl16*, *Cybb*, and *Csf1*) and downregulated (*P2ry12*, *Rhob*, *Nrip1*, *Cxxc5*, *Il21r,* and *Tgfbrap1*) in female 5xFAD microglia compared to their male counterparts (S.Fig.4f, g). Taken together, these data show that microglia exhibit sexually distinct transcriptomic profiles in the 5xFAD brain with female microglia engaging in ribosome biogenesis and glycolytic metabolism to a greater extent compared to male microglia.

### Interferon-stimulated genes are enriched in female 5xFAD microglia compared to males

Interferon-responsive microglia (IRM) have been identified near plaques in AD human and mouse model brains^15,68–72^. IRMs are characterized by the expression of genes in the *Ifit* (*e.g*., *Ifit3*, *Ifitm3*, *Ifit1*) and *Oasl* family (*e.g*., *Oasl2*), and include certain transcription factors (*e.g*., *Irf7*, *Irf3*, *Stat1*)^50^. Interestingly, we found that IRM genes were enriched in female 5xFAD cortical microglia compared to their male counterparts (Fig.5). We utilized a previously compiled list of interferon-stimulated genes (ISGs) ^68,73^ to further investigate how ISGs were present in female 5xFAD microglia compared to male counterparts. This list includes ISGs that have been identified in the CNS and are expressed by neurons, astrocytes, and microglia^68,73^. We found that 13% of all DEGs between female and male 5xFAD microglia were ISGs (60 ISGs out of 439 DEGs; Fig.5a). A heatmap of the expression levels of ISGs revealed stark sex differences with all 60 ISGs being upregulated in female 5xFAD microglia compared to male counterparts (Fig.5b). These genes are involved in the type I and type II IFN response^74^ and include IFN pathway initiation genes such as *Irf7, Stat1,* and *Stat2,* as well as canonical ISGs such as *Mx1, Ifitm3, Ifit3b, Ifit3, Ifi204, Ifi209, Isg15, Oas1g,* and *Oasl2*, amongst others (Fig.5b). Interestingly, *Usp18*, a negative regulator of IFN signaling^75^, was also upregulated in female 5xFAD microglia (Table S2), suggesting that microglia are attempting to contain this exacerbated response via a negative feedback loop. Apart from the canonical IFN-related genes, antigen presentation genes (*H2-D1*, *H2-K1*, *H2-Q5*, *H2-Q6*, *H2-T-ps*, *H2-T23*, *Psmb9*, *Psme1*, *Fcgr4*, and *Tap1*) which have also been identified to be ISGs^68,73^, were upregulated in female 5xFAD microglia, with the exception of Y chromosome-linked *Kdm5d* (Fig. 5b, S.Fig.4h-j). GO analysis revealed that female 5xFAD microglia exhibited an enrichment in pathways such as “response to type-I interferon”, “cellular response to interferon-beta”, “response to interferon-alpha”, and “response to type-II interferon”, as well as “response to virus” (Fig.4f; Table S3), a cellular process which involves IFN signaling and ISG expression^74^. In some instances, 5xFAD male microglia did not upregulate IFN genes (*Isg15*, *Irf7*, *Oas1g*; Fig.5c) compared to WTs, while female 5xFAD microglia did, suggesting that male microglia may exhibit a blunted IFN response to Aβ plaques at 5.5 months of age. These results suggest that IFN signaling is sexually distinct, with female 5xFAD microglia having a greater IFN response compared to their male counterparts. Collectively, these results further reveal sex differences in the microglia transcriptomic profile in response to Aβ, that is independent of estrous cycle stage.

**Figure 5.**
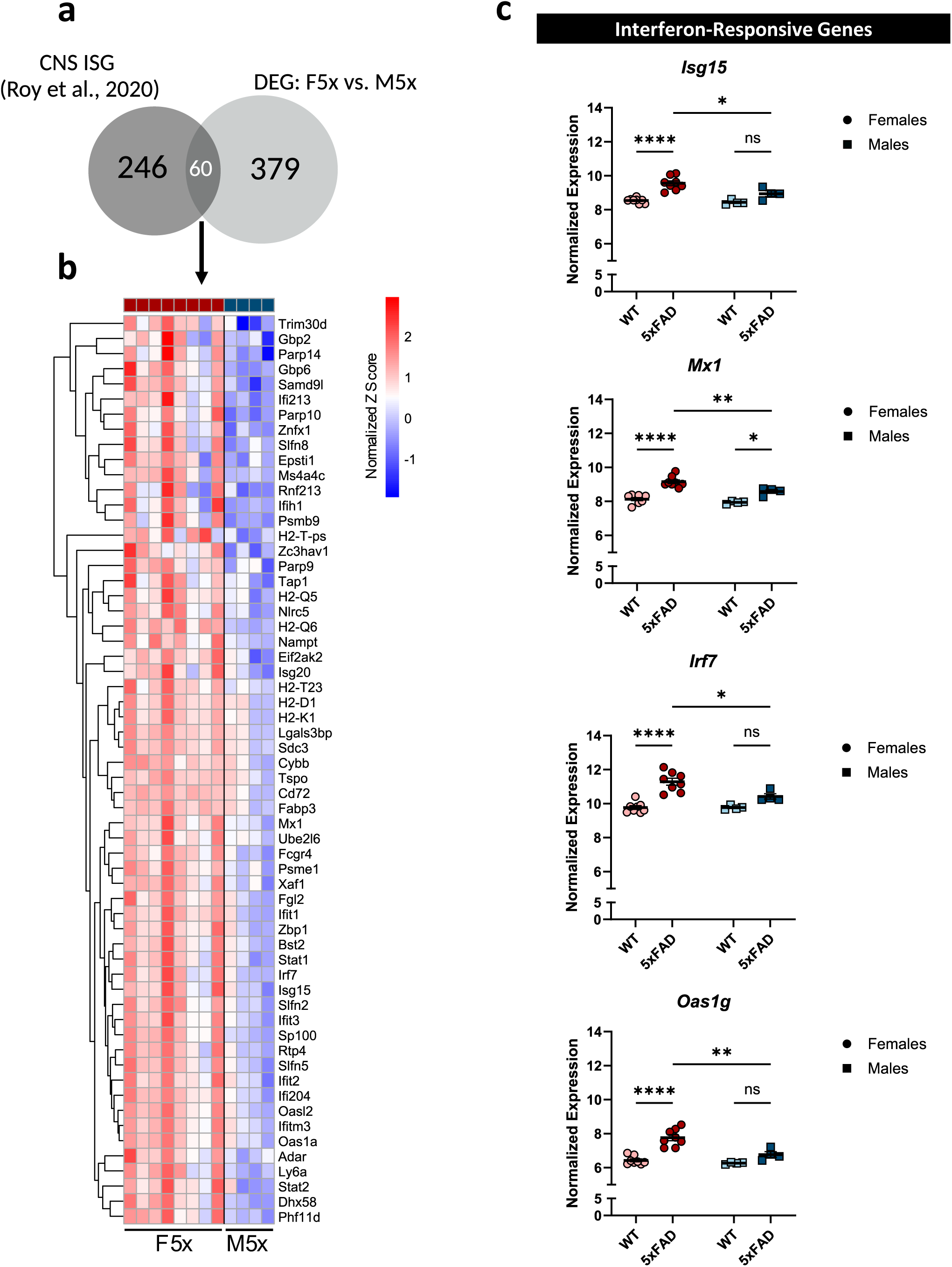
Female cortical microglia upregulate numerous interferon-stimulated genes in response to amyloid-β. a) Venn diagram of all the DEGs (including up- and downregulated) resulting from the pairwise comparison between female and male 5xFAD microglia that are shared with previously reported CNS ISGs. F5x = female 5xFAD microglia, M5x = male 5xFAD microglia. b) Heatmap of the 60 differentially expressed ISGs from the pairwise comparison between female and male 5xFAD microglia as identified in (a). c) Transcript levels (rLog Transformed FPKM) of *Isg15*, *Mx1*, *Irf7,* and *Oas1g* in female and male WT and 5xFAD microglia. Two-way ANOVA and Tukey’s post hoc test; *\*p < 0.05, **p <0.01, ***p<0.001, ****p<0.0001.* Data is presented as mean ± SEM.

## Discussion

Understanding how sex contributes to disease pathogenesis, progression, and prognosis is pivotal to developing personalized disease-modifying therapies. In this study, we assessed the sex differences in Aβ plaque morphology and microglia-plaque interaction, as well as investigated sex and estrous cycle stage differences in microglia transcriptomic profiles in the 5xFAD mouse model of amyloidosis. We found that female 5xFAD brain cortices had increased fibrillar Aβ plaque volume and decreased plaque sphericity, a measure of plaque compaction. We also found that, although there was no difference in microglia-plaque coverage, female microglia encapsulated plaques with low compaction to a greater extent compared to male microglia. When assessing the microglial transcriptome in the brains in which Aβ was or was not present, we found no overt differences between microglia at two different stages of the estrous cycle. Likewise, female and male WT microglia did not substantially differ, except for the sex-specific expression of sex chromosome-linked genes. However, female and male 5xFAD microglia differed in more than just sex-chromosome linked genes, as the majority of sexually differentiated genes were autosomal genes involved in glycolysis, antigen presentation, DAM/MGnD, and IFN signaling. Taken together, this study reveals that sex, and not estrous cycle stage, significantly affects the microglial transcriptome and that microglia are sexually differentiated in the context of AD.

### Sex differences in plaque morphology and microglia-plaque interaction

Various studies have reported that microglia may function as plaque “factories” leading to the initial seeding of Aβ plaques and contributing to their continuous growth^19,23,76^. This is supported by pharmacological and genetic approaches that deplete microglia which reveal that plaques fail to form in the brain parenchyma if microglia are absent during the early stages of Aβ pathology^23,53,77–80^. It is postulated that microglia phagocytose Aβ species (*e.g.,* soluble and oligomeric conformations), which subsequently aggregate in the acidic lysosome, where the pH is optimal for Aβ fibrilization^17,20,76^. Longitudinal two-photon studies demonstrate that microglia then seed these plaques via exocytosis or through apoptosis^76^. Apart from their role in the initial seeding, microglia have also been implicated in promoting plaque growth via continuous deposition of phagocytosed Aβ onto existing plaques^19,23,76^. Other studies have noted that dense-core plaque deposition may be facilitated by TAM (TYRO3, AXL, MER) receptor-dependent microglia Aβ phagocytosis^19^. Our findings indicate that female microglia have enhanced Aβ phagocytosis, which could, on the basis of the mechanisms mentioned above, potentially explain the larger size of the female plaques we observed.

Apart from seeding and growing plaques, microglia also remodel plaque ultrastructure over time^12,13,53^. Herein, we presented various plaque morphological features that microglia are known to modulate. We found a difference in fibrillar plaque volume and sphericity (3D analyses) between female and male 5xFAD mice. This is consistent with a previous report that noted a sex difference in plaque area and circularity (2D analyses) in EFAD (5xFAD-ApoE3) mice^54^. Similarly to our results, this prior study reported greater plaque area and lower plaque circularity in female compared to male brains^54^. Interestingly, we found that microglia coverage of the plaques was not sexually differentiated, contrasting with findings by Stephen et al., 2019 who reported that males had greater microglia-plaque coverage compared to females. It is worth noting, however, that these observations could have been influenced by the presence of the human ApoE3 variant, which is not present in our model.

### Female and male 5xFAD microglia are sexually distinct, independently of estrous cycle stage

The relative contributions of sex chromosomes and gonadal hormones to disease pathogenesis has yet to be disentangled, particularly regarding CNS immune responses. Interestingly, a previous study showed that ventral hippocampal neurons differed across the proestrus and diestrus stages when assessing chromatin organization and transcriptomic profiles^43^. Whether other CNS cell types exhibit a similar fluctuation remains to be elucidated. Microglia express hormone receptors such as estrogen receptor α and β as well as G-coupled receptors (GPR30) making them capable of responding to these pleiotropic hormones^81^. Various studies have empirically tested this by administering exogenous steroids and observing a microglial response^82–84^. However, to our knowledge, no other study has investigated whether microglia respond to endogenous changes in the hormonal milieu in gonadally intact mice. To address this gap, we assessed two hormonally distinct stages of the rodent estrous cycle (proestrus and diestrus) that mimic the late follicular and luteal phase of the menstrual cycle in humans and exhibit high and low levels of estrogen, respectively^50^. Except for a few genes, we did not find extensive differences in the microglial transcriptome at different stages of the estrous cycle in WT or 5xFAD mice. Our results indicate that at 5.5 months of age, endogenous fluctuations in gonadal hormones do not overtly influence the female microglial transcriptome in the healthy brain or in the context of Aβ pathology in the 5xFAD mouse model. This is consistent with a recent study that collected peripheral monocytes from women at different stages of their menstrual cycle (follicular and menstrual stages) and differentiated them into macrophages *in vitro*^59^. The investigators did not observe any differences in the enrichment of inflammatory response genes linked to AD at these different stages, although they noted that the time spent *in vitro* could have impacted these results^59^. Notably, however, one of the genes that we found to be differentially expressed and that showed higher expression in proestrus female microglia compared to diestrus female microglia in both WT and 5xFAD mice was *Nlrp3*, a component of the inflammasome that is involved in the production of IL-1β and IL-18^85^. Interestingly, previous reports in humans have identified that monocular cells derived from women in the follicular (proestrus-like) phase of the menstrual cycle produce greater amounts of IL-1 species (IL-1α and IL-1β) compared to those isolated from women in the luteal (diestrus-like) phase in the absence of any stimulation^86^.

Previous studies investigating sex differences in microglia have mainly focused on the hippocampus, however, microglia are known to be regionally heterogeneous^28,87^. For this reason, we wanted to examine the female and male microglial transcriptome in another disease-relevant brain region: the cerebral cortex. We found that, similar to the hippocampus, female cortical microglia tended to shift their metabolism toward glycolysis in response to Aβ pathology^66,88^. Recent studies have found that this enhanced glycolytic metabolism seems to be more pronounced in aging WT microglia of both sexes, but particularly in older female microglia compared to older male microglia^63,88^. This suggests that sex divergence occurs as part of healthy aging and might contribute to sex differences in AD, given that age is the primary risk factor for developing this disease. Moreover, a sex difference in glycolytic gene expression and glycolytic functions was noted in another study using the APP-PS1 mouse model of amyloidosis, where female microglia had elevated levels of glycolytic genes^66^. Interestingly, the expression of these glycolytic genes seems to be prominent in microglia that are in close proximity to plaques and adopt a DAM signature as postulated in a study using the 5xFAD mouse model^88^. Although not as efficient as oxidative phosphorylation (OXPHOS) in producing high quantities of ATP, glycolysis produces ATP at a faster rate compared to OXPHOS^65^. It is thought that this metabolic reprogramming to glycolysis enables microglia to perform high energy-demanding functions such as phagocytosis^65^.

Apart from immunometabolic reprogramming, microglia are known to adopt specific transcriptional programs, namely DAM^14^ and MGnD^16^. These two phenotypes largely overlap and underscore the functional roles of microglia in the context of neurodegeneration^14,16^. For instance, the MGnD phenotype is seen in microglia that phagocytose apoptotic neurons and, like DAM, these cells are found in close proximity to plaques^16^. Our findings reveal that both DAM and MGnD genes are enriched in female compared to male 5xFAD microglia. Notably, *ApoE,* which is necessary for the MGnD program and is involved in lipid metabolism and transport^16^, was significantly increased in female 5xFAD microglia compared to their male counterparts. This could have important implications as *ApoE* confers the greatest genetic risk of developing late-onset AD with the *ApoE4* variant disproportionately increasing the risk of AD in women in a dose-dependent manner^89,90^. Interestingly, however, we did not find canonical markers of these DAM/MGnD phenotypes, such as *Trem2*, *Clec7a*, *Cst7, Lpl, Cd9, Spp1* and *Axl*, amongst others, to be sexually differentiated^14^. Notably, this set of genes belongs to stage 2 DAM (*Trem2*-dependent stage), whereas *ApoE*, *Fth1*, *H2-D1*, and *Lyz2*, which did show sex-specific expression, belong to stage 1 DAM (*Trem2*-independent stage)^14^, an observation that could be due to the age (*i.e*., 5.5 months of age) assessed in this study. Overall, our results suggest that female microglia have greater levels of DAM genes, consistent with previous reports.

In addition to these phenotypes, microglia are known to secrete cytokines that modulate the disease. IFNs are cytokines commonly secreted by immune cells, including microglia, in response to viral infections^74^. Interestingly, IFNs seem to play a role in AD pathogenesis, but their exact function is currently understudied. Some reports have suggested that IFITM3, a prominent ISG, enhances γ-secretase activity, leading to more Aβ production^91^. Furthermore, a specific microglial transcriptomic signature characterized by IFN response genes was identified in AD post-mortem brains and AD mouse models^59,68^. This IRM signature is characterized by the expression of ISGs (*Ifitm3*, *Isg15*, *Mx1*, *Oasl2*) as well as IFN-related genes (*Stat1*, *Stat2*, *Irf7*), and microglia seem to adopt this profile when proximal to Aβ plaques^68–70^. Our results show that female 5xFAD microglia have increased levels of IFN genes compared to their male counterparts (Fig.4, 5). Whether this increase in IFN expression is due to greater Aβ burden or enhanced Aβ phagocytosis by female microglia remains to be dissected. One hypothesis supports the latter and argues that increased IFN expression and secretion may be associated with the internalization of Aβ plaques that contain potent stimulators of the IFN pathway (e.g., nucleic acids such as DNA and RNA)^68^. However, one could also argue that phagocytosis of cellular debris, which also contains nucleic acids, could similarly induce an IFN response.

Notwithstanding, this study has several limitations. For instance, we mainly present transcriptomic data and these results require validation at the protein level. We also use the 5xFAD mouse model of amyloidosis, which has been reported to contain a putative estrogen receptor element (ERE), a DNA motif where the estrogen receptor binds, upstream of the transgene promoter, Thy1^92^. However, as noted by the authors the ERE sequence contains a base pair mutation that deviates from the consensus sequence and thus is unclear whether it is active or even functional^92^. Furthermore, ERE presence does not dictate whether they may have a positive or negative impact on transcription and oftentimes multiple EREs are required to exert a functional impact^93^. Studies that report a sex bias in transgene expression only observe a modest sex difference in the *hAPP*, but not *hPSEN1* transgenes, both of which are under the same promoter, with some later studies not replicating the sex difference in *hAPP* transgene levels^92,94,95^. Our data suggests that there is not statistical difference in *hAPP* nor *hPSEN1* transgene expression in the 5xFAD mouse model, suggesting that the differences described here are not an artifact of the transgenic mouse model. Importantly, the sex differences in the microglial transcriptome we observed could be driven by the sex differences in Aβ burden rather than inherent sex-specific microglial phenotypes. Further correlational and single cell RNAseq studies should be performed at different ages in animals of both sexes to determine how microglial phenotype relates to amyloid pathology. Finally, we only report on proestrus and diestrus stages and do not include estrus or metestrus stages, for which we cannot exclude the possibility that microglia may differ at these other stages of the estrous cycle.

Overall, our study highlights the importance of including sex as a biological variable, especially in the context of disease. We present evidence that microglia are sexually differentiated in AD, making them potential targets for precision-based therapeutic interventions. We also provide evidence that endogenous fluctuations in hormones do not overly affect the microglial transcriptome in the healthy brain or in an amyloidosis model. However, we do highlight stark differences in transcriptional programs between the sexes, where female microglia transition to glycolytic metabolism, enhance DAM/MGnD, and IFN signaling, and male microglia retain a more homeostatic signature. Overall, whether these sex differences are cell-intrinsic, and microglia are divers of the sex differences in AD, or whether microglial phenotypes are driven by sexually distinct pathology remains to be unraveled.

## Supporting information

Supplemental Table 1. DAM genes in F5x v FWt and M5x v MWt

Supplemental Table 2. DAM genes in F5x v M5x

Supplemental Table 3. GO Results of F5x v M5x Upregulated Pathways

## Acknowledgements

We thank Mark Osabutey, Lee Trojanzcyk, Elizabeth Plunk, and Kirk Persaud for assistance with tissue procurement. We thank Dr. Olga Astapova, Department of Medicine, Endocrine/Metabolism, for her advice and training on vaginal smears and estrus cycle characterization. We thank Nikita Noyes-Martel, an undergraduate at the University of Rhode Island who participated in a Summer Research program at the University of Rochester for vaginal smear collection. We thank Dr. Linh Le for the design of the MeX04 flow cytometry experiment and Cassandra Lamantia for the MeX04 injections. We thank the Center for Advanced Light Microscopy and Nanoscopy (RRID: SCR_023177) and a special thank you to Dr. Kaye Thomas, Julie Zhang, and Dr. Emma Noris for their help with image acquisition/IMARIS and consultation. We thank the URMC Flow Cytometry Core (RRID: SCR_012360) for aiding in the flow cytometry/FACS experiments and consultation. We thank the Genomics Research Center (RRID: SCR_012359) for their help with the RNAseq experiment and consultation.

## Author contributions

LCR conceived the study in consultation with MKO. LCR designed, conducted, and carried out data analyses for most of the experiments in this study with the following contributions from co-authors: NNM, performed uterus weight dissections and assisted with vaginal smear collection/analysis and FACS experiment. JLB performed the RNAseq quality control, sequence alignment, rLog normalization of sequence reads/count, as well as produced the PCA plot and Euclidian heatmap. AKM provided suggestions on analysis of the morphological plaque data. LCR, AKM, and MKO wrote the first draft of the manuscript. All authors reviewed and contributed to the final version.

## Data availability

The data from this study are available from the corresponding author upon request.

---

**Supplemental Figure 1.**
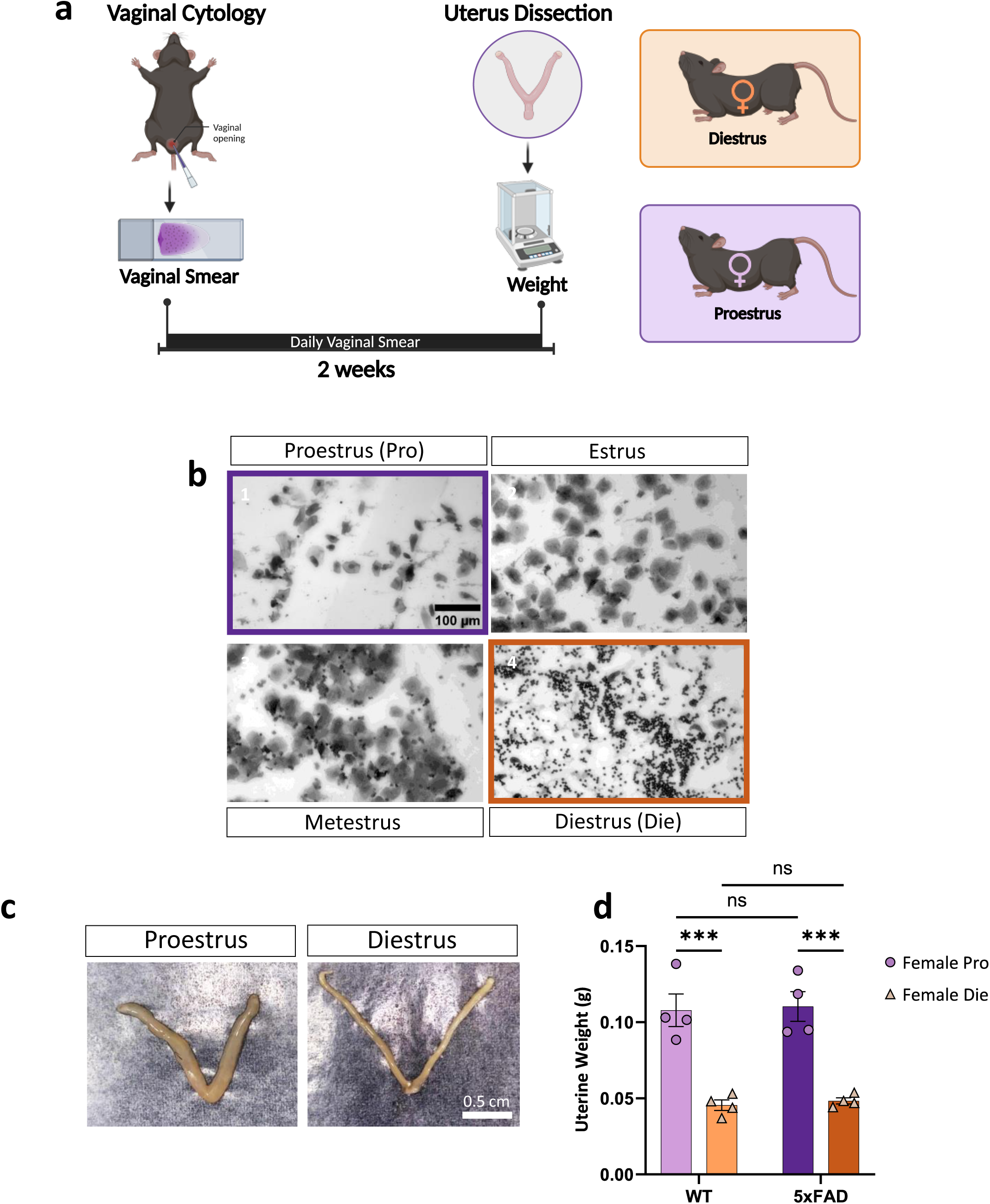
Vaginal cytology and uterine weight confirm WT and 5xFAD females cycle regularly. a) Schematic of experimental design: Vaginal smears were taken throughout a period of 2 weeks for cytological analyses and estrous cycle staging. Uterus weight at time of sacrifice was used to confirm correct staging into proestrus or diestrus stages. b) Representative images of vaginal cell composition at proestrus, estrus, metestrus, and diestrus stages (Scale bar = 100 µm). c) Representative image of dissected uteruses from a proestrus and a diestrus female (Scale bar = 1 cm). d) Uterus weight (g) of the proestrus and diestrus WT and 5xFAD females used in the RNAseq experiment. Two-way ANOVA and Tukey’s post hoc test; *p = 0.0004*. Data is presented as mean ± SEM.

**Supplemental Figure 2.**
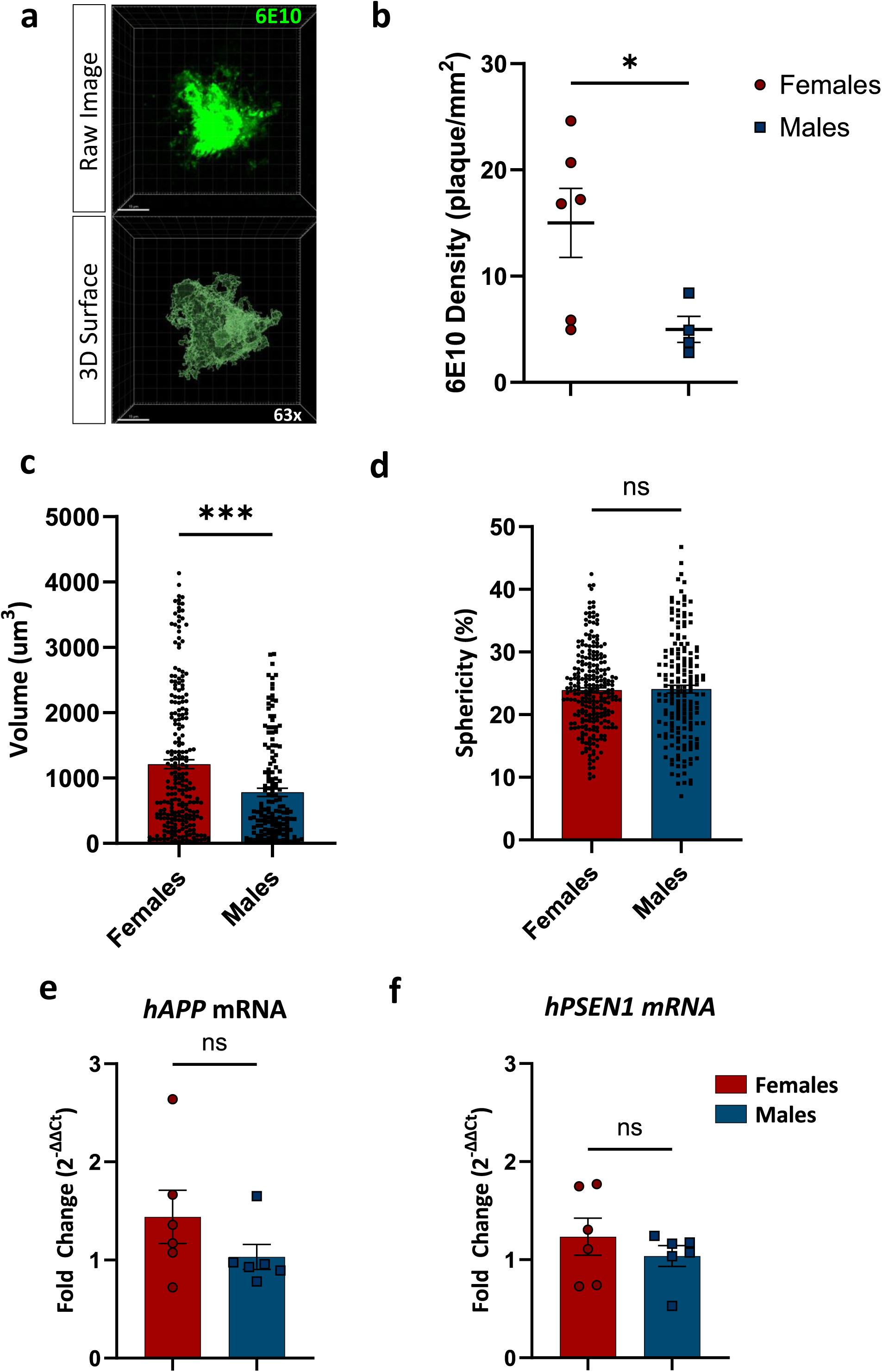
Sex differences in 6E10+ plaque density, plaque volume, but not plaque sphericity. a) Representative 63x confocal image (top) and 3D surface rendition (bottom) of 6E10+ plaque (Scale bar = 10 µm). b) Quantification of 6E10+ cortical plaque density (plaques/µm^2^) at 10x magnification for global plaque number assessment (n = 4-6 5xFAD mice/group). c) Quantification of cortical 6E10+ plaques volume (µm^3^) in female and male 5xFAD mice at 63x magnification. Each data point represents an individual plaque analyzed at high magnification (n = 152-237 plaques/group) derived from three 5xFAD mice/group. Two-tailed Mann Whitney test, *\*\*\*p=0.0001*. Data is presented as mean ± SEM. d) Quantification of cortical 6E10+ plaques sphericity (%) in female and male 5xFAD mice. Each data point represents an individual plaque analyzed at high magnification (n = 174-255 plaques/group) derived from three 5xFAD mice/group. Two-tailed Unpaired t test. Data is presented as mean ± SEM. e) mRNA quantification of human APP transgene in female and male 5xFAD mice (n = 6 5xFAD mice/group). Two-tailed Mann Whitney test (*p = 0.1797*). Data is presented as mean ± SEM. f) mRNA quantification of human PSEN1 transgene in female and male 5xFAD mice (n = 6 5xFAD mice/group). Two-tailed Mann Whitney test (*p = 0.4848*). Data is presented as mean ± SEM.

**Supplemental Figure 3.**
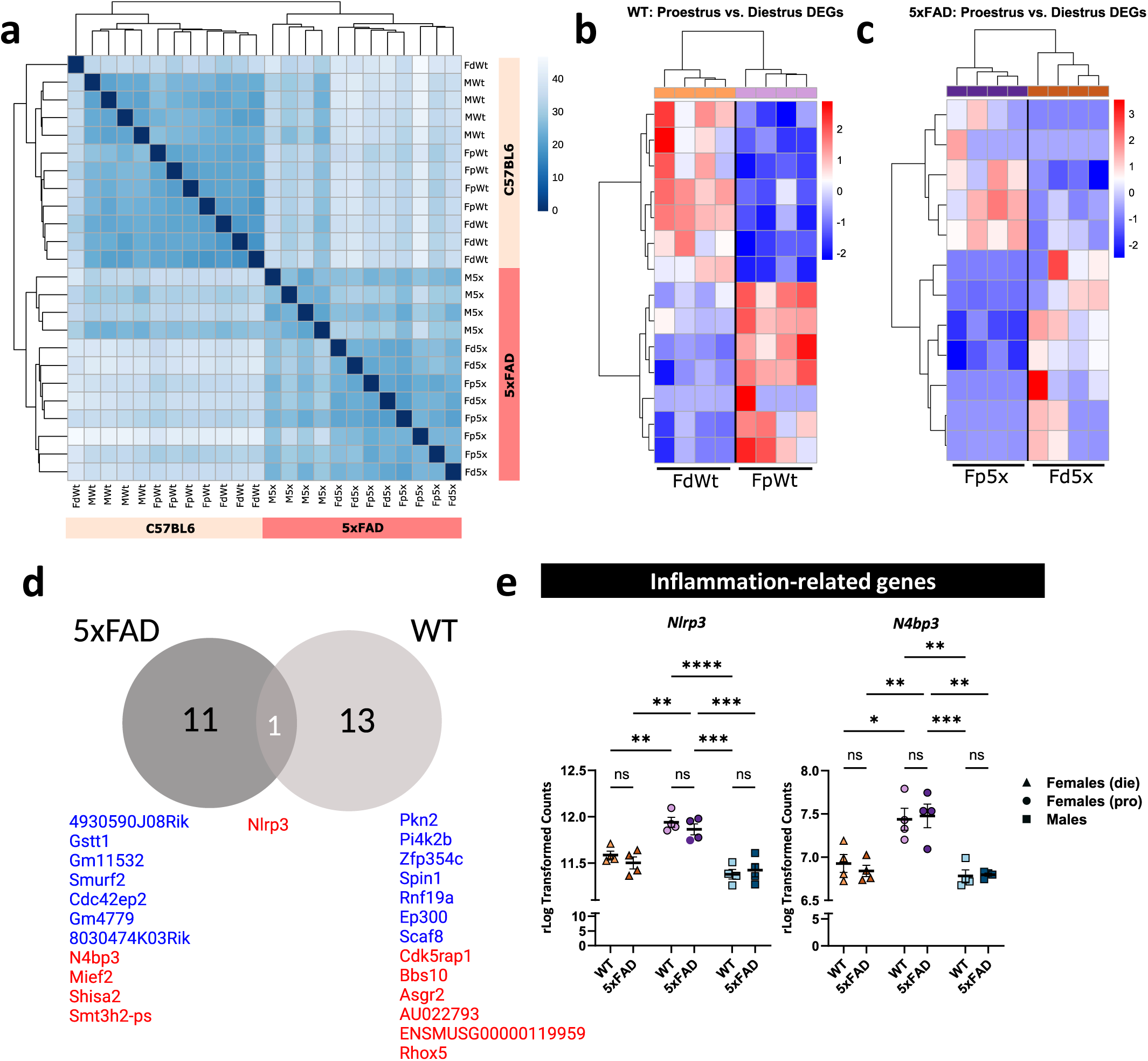
No overt differences in diestrus and proestrus female microglial transcriptome in WT nor 5xFAD mice. a) Euclidian heatmap showing the sample-to-sample distance, with less distance (*i.e.,* 0 or dark blue) meaning the samples are very similar to one another. b-c) Heatmap of the DEGs between proestrus and diestrus WT (b) and 5xFAD (c) microglia. d) Venn diagram of the DEGs between proestrus and diestrus female microglia shared across genotypes (5xFAD vs. WT) with a list of the genes that are upregulated (red) and downregulated (blue). e) Transcript levels (rLog Transformed FPKM) of *Nlrp3* and *N4bp3* in proestrus female, diestrus female, and male WT and 5xFAD microglia. Two-way ANOVA and Tukey’s post hoc test; *\*p < 0.02, **p <0.005, ***p<0.001, ****p<0.0001.* Data is presented as mean ± SEM.

**Supplemental Figure 4.**
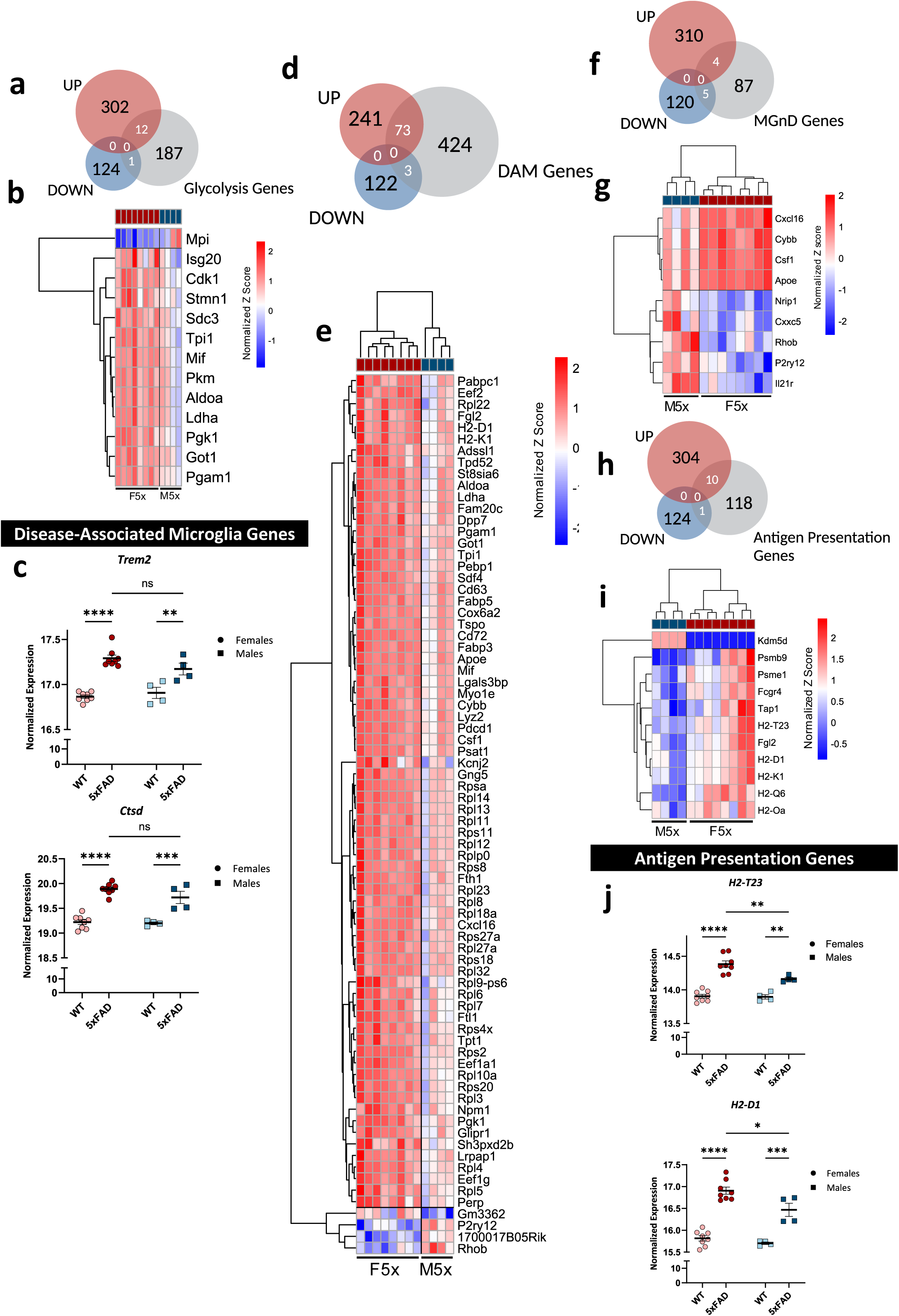
Female cortical microglia upregulate genes involved in glycolytic metabolism, DAM/MGnD phenotype, and antigen presentation in response to amyloid-β. a) Venn diagram of the DEGs (UP = upregulated, DOWN = downregulated) resulting from the pairwise comparison between female and male 5xFAD microglia that are shared with previously reported glycolysis genes. b) Heatmap of the 12 upregulated and 1 downregulated glycolysis genes from the pairwise comparison between female and male 5xFAD microglia as identified in (a). c) Transcript levels (rLog Transformed FPKM) of *Trem2* and *Ctsd* in female and male WT and 5xFAD microglia. Two-way ANOVA and Tukey’s post hoc test; *\*\*\*\*p<0.0001.* Data is presented as mean ± SEM. d) Venn diagram of the DEGs (UP = upregulated, DOWN = downregulated) resulting from the pairwise comparison between female and male 5xFAD microglia that are shared with previously reported DAM genes. e) Heatmap of the 73 upregulated and 3 downregulated DAM genes from the pairwise comparison between female and male 5xFAD microglia as identified in (d). f) Venn diagram of the DEGs (UP = upregulated, DOWN = downregulated) resulting from the pairwise comparison between female and male 5xFAD microglia that are shared with previously reported MGnD genes. g) Heatmap of the 4 upregulated and 5 downregulated MGnD genes from the pairwise comparison between female and male 5xFAD microglia as identified in (f). h) Venn diagram of the DEGs (UP = upregulated, DOWN = downregulated) resulting from the pairwise comparison between female and male 5xFAD microglia that are shared with previously reported antigen presentation genes. i) Heatmap of the 10 upregulated and 1 downregulated antigen presentation genes from the pairwise comparison between female and male 5xFAD microglia as identified in (h). j) Transcript levels (rLog Transformed FPKM) of *H2-T23* and *H2-D1* in female and male WT and 5xFAD microglia. Two-way ANOVA and Tukey’s post hoc test; *\*p < 0.05, **p <0.01, ****p<0.0001.* Data is presented as mean ± SEM.

**Supplemental Figure 5.**
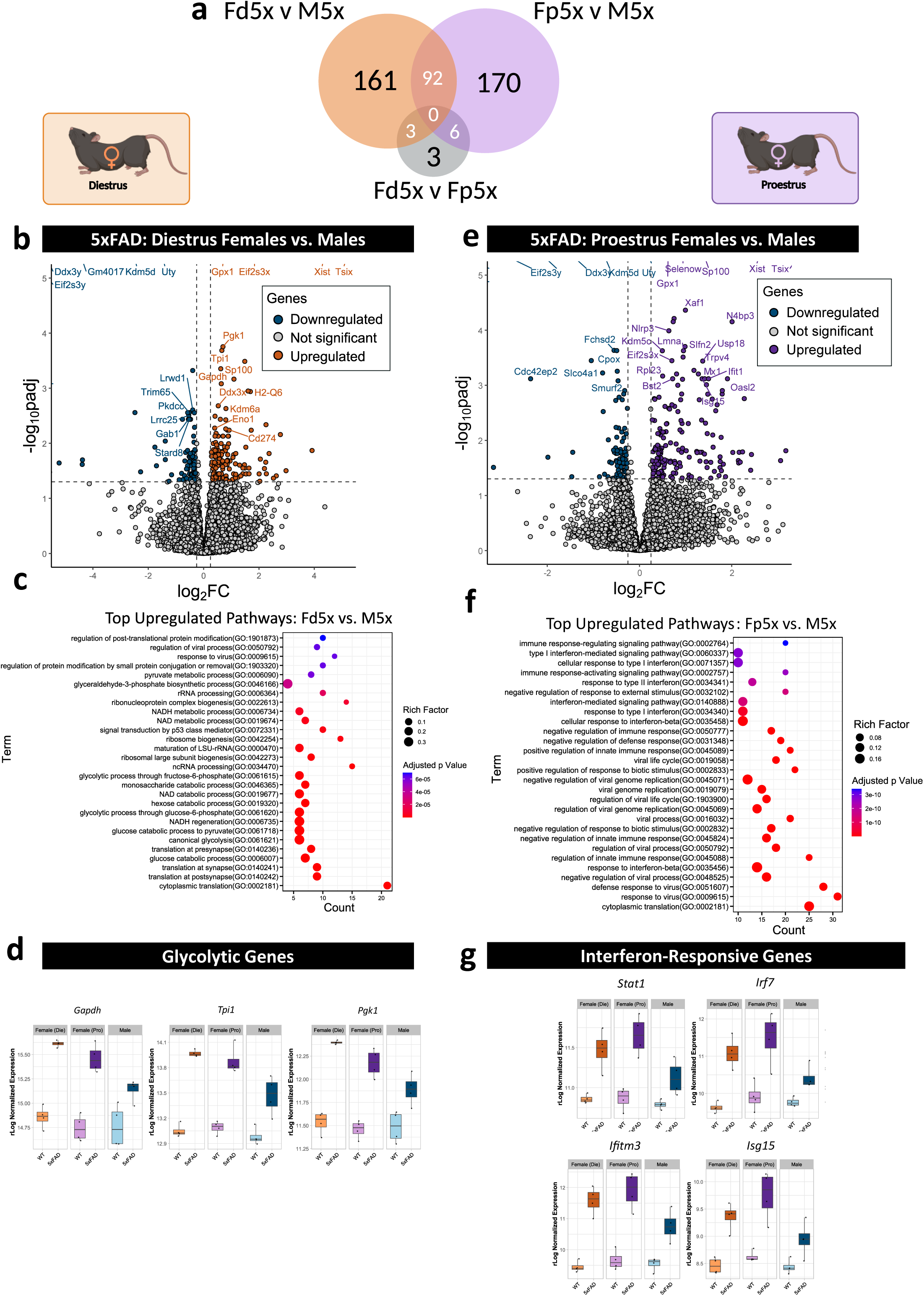
Proestrus and diestrus microglia engage distinct functional pathways in response to amyloid-β when compared to male microglia. a) Venn diagram of all the DEGs (including up- and downregulated) shared across various pairwise comparisons. Fd5x = diestrus female 5xFAD microglia, Fp5x = proestrus female 5xFAD microglia, M5x = male 5xFAD microglia. b) Volcano plot showing the DEGs in the pairwise comparison between diestrus female and male 5xFAD microglia. *FDR<0.05; |LFC|>0.25*. c) Gene ontology analysis of the DEGs from the pairwise comparison between diestrus female and male 5xFAD microglia. d) Boxplots showing the transcript levels (rLog Transformed FPKM) of key glycolytic genes (*Gapdh*, *Tpi*, *Pgk1*) that are differentially expressed between diestrus female and male 5xFAD microglia. Values shown for all sexes and genotypes. Data is presented as median ± IQR. e) Volcano plot showing the DEGs in the pairwise comparison between proestrus female and male 5xFAD microglia. *FDR<0.05; |LFC|>0.25*. f) Gene ontology analysis of the DEGs from the pairwise comparison between proestrus female and male 5xFAD microglia. g) Boxplots showing the transcript levels (rLog Transformed FPKM) of key interferon-related genes (*Stat1*, *Irf7*, *Ifitm3*, *Isg15*) that are differentially expressed between proestrus female and male 5xFAD microglia. Values shown for all sexes and genotypes. Data is presented as median ± IQR.

